# Insights into animal septins using recombinant human septin octamers with distinct SEPT9 isoforms

**DOI:** 10.1101/2021.01.21.427698

**Authors:** Francois Iv, Carla Silva Martins, Gerard Castro-Linares, Cyntia Taveneau, Pascale Barbier, Pascal Verdier-Pinard, Luc Camoin, Stéphane Audebert, Feng-Ching Tsai, Laurie Ramond, Alex Llewellyn, Mayssa Belhabib, Koyomi Nakazawa, Aurélie Di Cicco, Renaud Vincentelli, Jerome Wenger, Stéphanie Cabantous, Gijsje H. Koenderink, Aurélie Bertin, Manos Mavrakis

## Abstract

Septin GTP-binding proteins contribute essential biological functions that range from the establishment of cell polarity to animal tissue morphogenesis. Human septins in cells form hetero-octameric septin complexes containing the ubiquitously expressed SEPT9. Despite the established role of SEPT9 in mammalian development and human pathophysiology, biochemical and biophysical studies have relied on monomeric SEPT9 thus not recapitulating its native assembly into hetero-octameric complexes. We established a protocol that enabled the first-time isolation of recombinant human septin octamers containing distinct SEPT9 isoforms. A combination of biochemical and biophysical assays confirmed the octameric nature of the isolated complexes in solution. Reconstitution studies showed that octamers with either a long or a short SEPT9 isoform form filament assemblies, and can directly bind and cross-link actin filaments, raising the possibility that septin-decorated actin structures in cells reflect direct actin-septin interactions. Recombinant SEPT9-containing octamers will make it possible to design cell-free assays to dissect the complex interactions of septins with cell membranes and the actin/microtubule cytoskeleton.

**Summary:** Human septins in cells form hetero-octameric complexes containing the ubiquitously expressed SEPT9. Iv et al. describe the first-time isolation of recombinant human septin octamers with distinct SEPT9 isoforms. Reconstitution studies show that octamers with either a long or a short SEPT9 isoform form higher-order filament assemblies and directly bind and cross-link actin filaments.

## Introduction

Septins constitute a family of GTP-binding proteins conserved from algae and protists to mammals (Cao et al., 2007; Momany et al., 2008; Nishihama et al., 2011; Pan et al., 2007). Septins are involved in a wide range of biological processes, from the establishment of cell polarity and cell division to cell-cell adhesion, cell motility, animal tissue morphogenesis and infection (Fung et al., 2014; Marquardt et al., 2019; Mostowy and Cossart, 2012; Weirich et al., 2008). In human pathophysiology, a role of septins has been established in neuropathies, infertility and tumorigenesis (Dolat et al., 2014a; Montagna et al., 2015). Despite their essential roles, how human septins organize and function in cells remains much more poorly understood than for budding yeast, in which septins were first discovered (Hartwell, 1971; Hartwell et al., 1970). Mammalian septins are thought to associate with cell membranes (Akil et al., 2016; Bridges et al., 2016; Damalio et al., 2013; Dolat and Spiliotis, 2016; Omrane et al., 2019; Tanaka-Takiguchi et al., 2009; Zhang et al., 1999) like their yeast counterparts (Bertin et al., 2010; Bridges et al., 2016; Bridges et al., 2014; Casamayor and Snyder, 2003). Unlike budding yeast septins, mammalian septins localize extensively to actin and microtubules in cells, for example to the ingressing cytokinetic ring in dividing cells (Estey et al., 2010; Joo et al., 2007; Kim et al., 2011; Kinoshita et al., 1997; Surka et al., 2002), stress fibres in interphase cells (Calvo et al., 2015; Connolly et al., 2011; Dolat et al., 2014b; Joo et al., 2007; Kim et al., 2011; Kinoshita et al., 2002; Kinoshita et al., 1997; Liu et al., 2014; Surka et al., 2002; Verdier-Pinard et al., 2017; Xie et al., 1999; Zhang et al., 1999), and to interphase, mitotic spindle, and intercellular bridge microtubules (Bowen et al., 2011; Nagata et al., 2004; Nagata et al., 2003; Spiliotis et al., 2008; Spiliotis et al., 2005; Surka et al., 2002; Verdier-Pinard et al., 2017). Mammalian septin association with membranes as well as with the actin and microtubule cytoskeleton has made it difficult to dissect how they function, and at the same time raises the intriguing possibility that septins mediate cytoskeleton-membrane cross-talk.

Studies of native and recombinant septins isolated from budding yeast (Bertin et al., 2008; Farkasovsky et al., 2005; Frazier et al., 1998; Garcia et al., 2011; Versele and Thorner, 2004), *Drosophila* (Field et al., 1996; Huijbregts et al., 2009; Mavrakis et al., 2014), *C. elegans* (John et al., 2007), and mammalian cell lines and tissues (Hsu et al., 1998; Kim et al., 2011; Kinoshita et al., 2002; Sellin et al., 2011; Sirajuddin et al., 2007) have established that septins form heteromeric complexes, with each septin present in two copies, forming a palindrome. Phylogenetic analysis has classified human septins in four homology groups, namely the SEPT2 group (SEPT1, 2, 4, and 5), SEPT6 group (SEPT6, 8, 10, 11, and 14), SEPT7 group (SEPT7), and SEPT3 group (SEPT3, 9, and 12) (Kinoshita, 2003) (see Materials and methods for nomenclature). Native human septins isolated from cells exist in the form of stable hexamers and octamers (Kim et al., 2011; Sellin et al., 2011; Sellin et al., 2014). Hexamers are composed of septins from the SEPT2, SEPT6, SEPT7 groups, while octamers contain additional septins from the SEPT3 group (Fig. 1 A).

**Figure 1.**
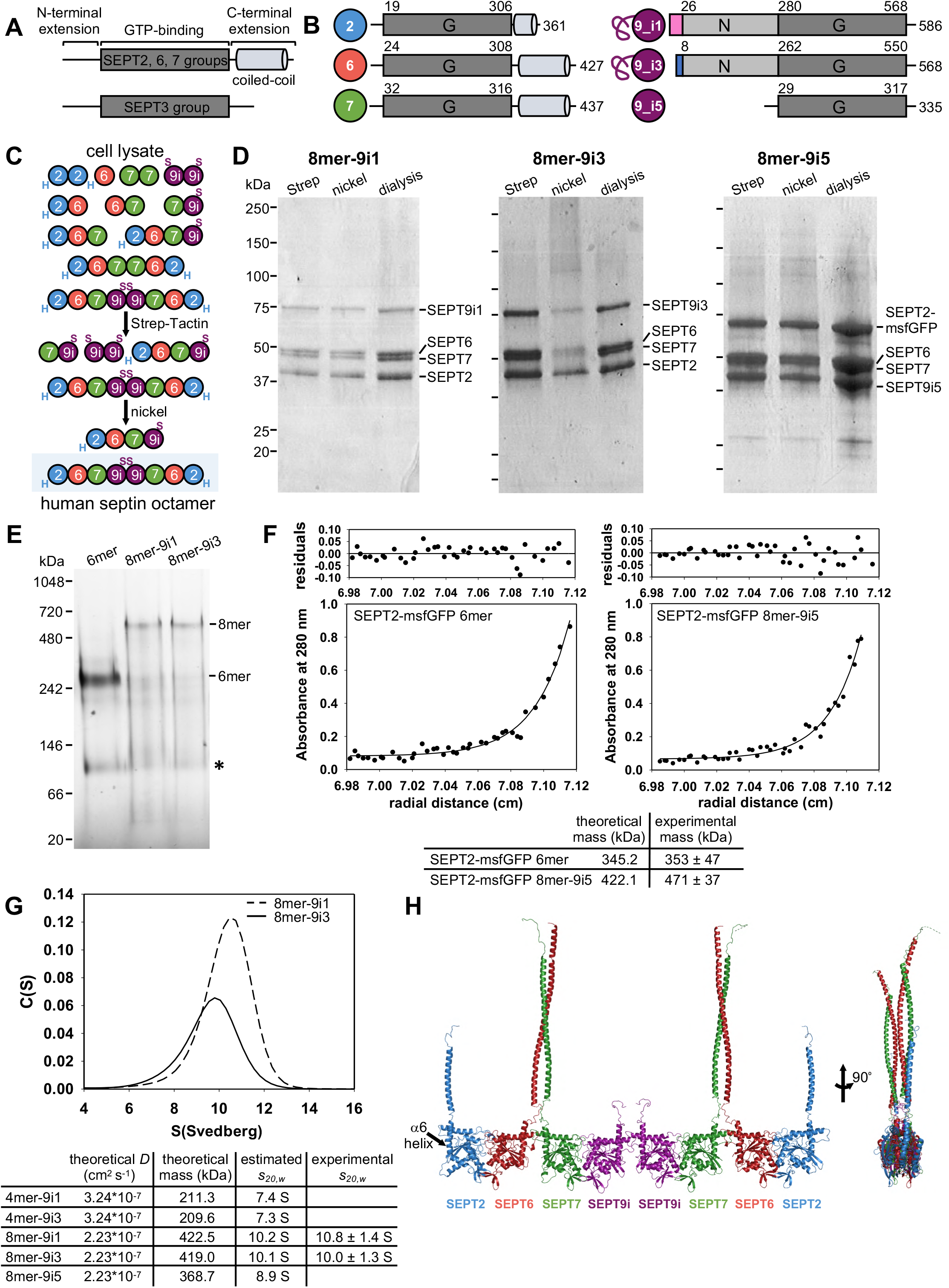
Isolation and characterization of recombinant human octamers containing distinct SEPT9 isoforms. **(A-B)** Schematic representation of a mammalian septin showing the conserved GTP-binding (G) domain flanked by N- and C-terminal extensions. There is experimental evidence that the N-terminal extension, at least for some septins, is intrinsically disordered (Garcia et al., 2006). With the exception of SEPT3 group septins, the C-terminal extension is predicted to contain a coiled-coil (A). Schematics in (B) depict the human septins used in this study, their size indicated by the very C-terminal residue number. Residue numbers at the start and end of the G-domains correspond to the start of the α0 helices and the end of the α6 helices, respectively. Residue numbers right after the isoform-specific sequences for SEPT9_i1 and SEPT9_i3 indicate the start of their shared long N-terminal extension. The last 28 residues of this extension constitute the N-terminal extension of the short isoform, SEPT9_i5. Color-coded spheres depicting the different subunits throughout the manuscript are shown next to the respective septins. The freehand line preceding the SEPT9 G domain of the long isoforms depicts their large N-terminal extension. **(C)** Schematic overview of the two-tag affinity purification scheme for isolating stoichiometric SEPT9-containing octamers. Upon septin co-expression in the bacteria cytoplasm, septins are expected to form stable hexamers and octamers (Kim et al., 2011; Sellin et al., 2011; Sellin et al., 2014). Other hypothetical homo- and hetero-subcomplexes could also form (Kim et al., 2012; Rosa et al., 2020; Valadares et al., 2017). A first Strep-tag affinity column isolates all Strep-tagged SEPT9 complexes (“S” for Strep-tag). A second nickel affinity step further isolates His_6_-tagged SEPT2-containing complexes (“H” for His_6_-tag), thus purifying SEPT2-SEPT6-SEPT7-SEPT9 complexes. **(D)** SDS-PAGE analysis of the purification of human septin octamers containing SEPT9_i1 (left, 8mer-9_i1), SEPT9_i3 (middle, 8mer-9_i3) and SEPT9_i5 (right, 8mer-9_i5). Coomassie-stained gels show fractions eluting from the Strep-tag affinity column, from the nickel affinity column and after the final dialysis step. Molecular weight markers are shown on the left; the same markers were used in all gels. The 8mer-9_i1 and 8mer-9_i3 complexes shown are nonfluorescent, whereas the 8mer-9_i5 complexes shown contain msfGFP-tagged SEPT2. The identification of the bands corresponding to the different septins was based on mass spectrometry and Western blot analysis (Fig. S1 F). See Materials and methods for the theoretical and apparent molecular masses. **(E)** Purified recombinant hexamers (6mer), 8mer-9_i1 and 8mer-9_i3 analyzed by blue native PAGE, followed by Coomassie-staining. Molecular weight markers are shown on the left. The apparent molecular masses for the recombinant 6mer and 8mer-9_i1 and −9i_3 are in line with the molecular masses of native hexamers and octamers isolated from human cell lysates (Sellin et al., 2014) (see also Materials and methods). The asterisk points to the presence of presumed septin monomers/dimers (Sellin et al., 2014). **(F and G)** Analytical ultracentrifugation of recombinant septin complexes. Sedimentation equilibrium experiments (F) of 1.5 mg.mL^−1^ of 6mer (left) and 8mer-9_i5 (right) at 11,000 rpm and 4°C. The filled circle symbols show the experimental radial concentration distribution at sedimentation equilibrium and the solid lines represent the best fit curves with the single-ideal species model. The residuals representing the variation between the experimental data and those generated by the fit are shown above the respective curves. The obtained experimental molecular masses are indicated in the table below the curves. (G) shows the sedimentation coefficient distributions *c(s)* of 0.3 mg.mL^−1^ 8mer-9_i3 (dark line) and of 0.5 mg.mL^−1^ 8mer-9_i1 (dashed line) obtained from sedimentation velocity experiments at 40,000 rpm and 20°C. The experimental sedimentation coefficients *s_20,W_* obtained are indicated in the table below the curves. **(H)** Model of a human SEPT2-SEPT6-SEPT7-SEPT9-SEPT9-SEPT7-SEPT6-SEPT2 octamer built using coiled-coil- and homology-modelling software (see Materials and methods for details). An *en face* view (left) and a side view after a 90° rotation (right) are shown. Coiled-coils are shown to extend along the same axis as the α6 helix (arrow). The generated model was used to calculate the theoretical translational diffusion coefficient, *D* (cm^2^.s^−1^), and the latter to further calculate its theoretical sedimentation coefficient for comparison with the experimentally obtained one (table in G) (see Materials and methods for details).

A well-documented feature of septins is that purified septin heteromeric complexes self-assemble into filaments (Valadares et al., 2017). Whether all native septin pools are filamentous, and how septin function is linked to the relative distributions of hexamers and octamers and their polymerization capacity within cells are not known. The most convincing evidence that septins form filaments *in vivo*, and that septin function depends on their ability to assemble into filaments, comes from budding yeast (Bertin et al., 2012; Byers and Goetsch, 1976; McMurray et al., 2011; Ong et al., 2014; Rodal et al., 2005). A powerful tool for studying septin assembly and function has been the use of recombinant septin complexes. Earlier studies using recombinant mammalian septin complexes have combined septins from two or more species, most likely for pragmatic reasons. Mouse SEPT2 was combined with human SEPT6 and SEPT7 (Kinoshita et al., 2002; Mavrakis et al., 2014; Sirajuddin et al., 2007), or with human SEPT6, SEPT7 and SEPT3 (DeRose et al., 2020), and mouse SEPT2 was also combined with human SEPT6 and rat SEPT7 (Bai et al., 2013). Furthermore, the longest SEPT7 isoform (UNIPROTKB identifier Q1681-1) was originally annotated with its N-terminus missing 19 residues (Macara et al., 2002). There are currently no studies showing whether these specific species-related differences affect septin function. Still, taking into account that these differences lie in exposed residues in the very N- or/and C-terminal extensions (Fig. 1 A, B), or within exposed loops in the GTP-binding domain, and given how poorly we understand the factors that impact animal septin assembly and function, there is a clear need to produce septin complexes with full-length septins from one species, notably human septin octamers containing SEPT2, SEPT6, SEPT7, and SEPT9.

SEPT9 is the only septin from the SEPT3 group whose expression is ubiquitous across human tissues, with SEPT3 and SEPT12 being neuron- and testis-specific, respectively (Cao et al., 2007; Connolly et al., 2011; Hall et al., 2005). *Sept9* gene knockout in mice is embryonic lethal (Fuchtbauer et al., 2011), and a large body of literature has implicated SEPT9 in diverse human cancers (Dolat et al., 2014a; Montagna et al., 2015). There are five SEPT9 isoforms (SEPT9_i) differing in the length and composition of the N-terminal extension preceding the GTP-binding domain (Connolly et al., 2014; McIlhatton et al., 2001) (Fig. 1 B). Distinct SEPT9 isoforms can have different functions, as reported for cytokinesis and cancer cell migration (Estey et al., 2010; Verdier-Pinard et al., 2017). Despite its importance in mammalian development and human pathophysiology, biochemical and biophysical studies of SEPT9 have been limited to the use of monomeric SEPT9 and fragments thereof (Bai et al., 2013; Dolat et al., 2014b; Nakos et al., 2019; Smith et al., 2015), thus not recapitulating its native assembly into hetero-octameric complexes (Sellin et al., 2011; Sellin et al., 2014). Multiple studies have documented promiscuity in septin-septin interactions in the absence of their physiologically relevant binding partners, affecting the availability of specific structural elements for interactions with other septins or interacting proteins (Castro et al., 2020; Valadares et al., 2017). The necessity to study septins in the context of their native heteromeric complexes is highlighted by the increasing number of structural studies of the factors governing the molecular specificity that determines the correct pairing of septins during complex assembly (Kumagai et al., 2019; Rosa et al., 2020; Sala et al., 2016).

The N-terminal extension in the long SEPT9 isoforms (SEPT9_i1, SEPT9_i2 and SEPT9_i3) is of considerable size (∼27-kDa, i.e. three-quarters of the size of the GTP-binding domain) making these isoforms the longest, in terms of the number of residues, of all human septins. Given that the long SEPT9 isoforms differ only in the composition of their N-terminal 25, 18 and 7 residues, respectively (Fig. 1 B), it is intriguing that they all associate with actin stress fibres in cells, whereas only SEPT9_i1 associates with microtubules (Nagata et al., 2004; Nagata et al., 2003; Surka et al., 2002). Different cell types express different sets of SEPT9 isoforms, with some cell types expressing specific long SEPT9 isoforms, and others lacking altogether long SEPT9 isoforms (Burrows et al., 2003; Sellin et al., 2014; Verdier-Pinard et al., 2017). Hereditary neuralgic amyotrophy (HNA), a rare neuropathy, has been mapped to missense mutations and duplications in the large N-terminal extension shared by the long SEPT9 isoforms (Collie et al., 2010; Hannibal et al., 2009; Kuhlenbaumer et al., 2005; Landsverk et al., 2009). Understanding SEPT9 function thus necessitates the isolation of recombinant septin octamers bearing distinct SEPT9 isoforms.

To enable studies of SEPT9 function in the context of its physiological assembly into hetero-octamers, we established a protocol that enabled, for the first time, the isolation of recombinant human septin octamers containing distinct SEPT9 isoforms (Fig. 1 B). A combination of biochemical and biophysical assays confirmed the octameric nature of the isolated octamers in solution, and also provided evidence for SEPT2 and SEPT9 occupying the end and central positions in the octamer. Fluorescence and electron microscopy showed that recombinant octamers containing either a long or a short SEPT9 isoform form higher-order filament assemblies. As a first step towards the reconstitution of recombinant SEPT9-containing octamers with known physiological interactors, we examined their interactions with actin filaments. Reconstitution studies showed that octamers with either a long or a short SEPT9 isoform directly bind and cross-link actin filaments, raising the possibility that septin-decorated actin bundles in cells reflect direct actin-septin interactions. Biochemical and biophysical reconstitution studies of recombinant octamers containing distinct SEPT9 isoforms with physiological septin interactors, such as membranes and microtubules, promise to provide a powerful complementary approach to cell and animal model studies of septin organization and function.

## Results and discussion

### A two-tag purification scheme yields stoichiometric recombinant human septin octamers containing distinct SEPT9 isoforms

To isolate octamers containing either a long SEPT9 isoform, in particular SEPT9_i1 and SEPT9_i3, or octamers containing a short SEPT9 isoform, SEPT9_i5 (Fig. 1 B), we combined the pET-MCN (pET <u>M</u>ulti-<u>C</u>loning and expressio<u>N</u>) series as a septin co-expression system (Diebold et al., 2011) with a two-tag purification scheme. We used two bicistronic vectors: one vector co-expressing SEPT2 and SEPT6, the other one SEPT7 and SEPT9_i (Fig. S1 A). To minimally perturb septin complex assembly, and interactions with other proteins or membranes, we chose small (1 kDa) tags, a hexahistidine (His_6_) tag and the eight amino-acid Strep-tag II (Fig. S1 B). To isolate octamers, we tagged the N-terminus of the end subunit, SEPT2, with a tobacco etch virus (TEV) protease-cleavable His_6_-tag, and the C-terminus of the central subunit, SEPT9_i, with a TEV-cleavable Strep-tag. The use of a Strep-Tactin affinity column to capture Strep-tagged SEPT9_i-containing complexes, followed by a nickel affinity column to retain the SEPT9_i-containing complexes that also bear His_6_-tagged SEPT2, is expected to isolate SEPT2-SEPT6-SEPT7-SEPT9 complexes (Fig. 1 C). We used this purification scheme to isolate both nonfluorescent septin complexes and fluorescent septin complexes containing SEPT2 with its C-terminus fused to monomeric superfolder GFP (msfGFP) (Costantini et al., 2012; Cranfill et al., 2016; Pedelacq et al., 2006; Zacharias et al., 2002). Indeed, SDS-PAGE analysis of the purification of human septin complexes containing SEPT9_i1 (octamers-9_i1), SEPT9_i3 (octamers-9_i3), or SEPT9_i5 (octamers-9_i5), followed by Coomassie staining, showed that our purification scheme succeeded to isolate SEPT2-SEPT6-SEPT7-SEPT9 complexes (Fig. 1 D). The assignment of the bands to the different septins was based on Western blot analysis and mass spectrometry. Western blots (Fig. S1 F), tryptic peptide coverage and pseudo-absolute quantitation of the mol fractions of proteins in our preps by mass spectrometry (Fig. S1 G and H) showed that the isolated complexes were >97% pure, intact and with SEPT2, SEPT6, SEPT7, SEPT9 in a 1:1:1:1 stoichiometry. We note that inverting the two columns, using batch affinity resins instead of prepacked columns, or combining prepacked columns and resins, all provided similar results (Fig. S 1 C-E).

Given that the purification scheme *per se* cannot distinguish between tetramers and octamers, we sought to determine if the isolated SEPT2-SEPT6-SEPT7-SEPT9 complexes were composed of tetramers or/and octamers by blue native PAGE followed by Coomassie staining (Fig. 1 E). For comparison, we included recombinant human SEPT2-, SEPT6-, SEPT7-containing hexamers that we isolated with the same purification protocol (Fig. S1 A). Blue native PAGE has been a powerful tool in detecting the presence and relative distributions of endogenous septin complexes in cell lysates, and is able to resolve septin tetramers from hexamers and octamers (Sellin et al., 2014). Native PAGE analysis of our hexamer and octamer preps showed bands that were in line with the presence of hexamers for the hexamer prep and octamers for the octamer preps, while providing no evidence for the presence of tetramers, suggesting that the latter either do not form, or they do so transiently. Our findings are consistent with SEPT9 being present in the form of stable octamers in cells (Sellin et al., 2011; Sellin et al., 2014).

To further corroborate the presence of a stable octameric population in our preps, we turned to analytical ultracentrifugation sedimentation equilibrium assays, comparing side by side hexamers and SEPT9_i5-containing octamers. Sedimentation equilibrium experiments provide an experimental measure of the absolute mass of proteins in solution (Taylor et al., 2015) and thus a powerful means of determining the species present in our preps. The obtained molecular masses from such experiments were consistent with the presence of hexamers for the hexamer preps, and the presence of octamers for the octamer preps, without any detectable evidence for tetrameric complexes in the octamer preps (Fig. 1 F). We complemented these assays with analytical ultracentrifugation sedimentation velocity experiments comparing octamer-9_i1 and octamer-9_i3 preps (Fig. 1 G). Sedimentation velocity assays measure the experimental sedimentation coefficient of proteins in solution and are thus able to detect the presence of multiple protein species. Sedimentation coefficients depend on the hydrodynamic properties of proteins, and are directly proportional to their mass and translational diffusion coefficient, the latter including the contribution of protein shape. To interpret the obtained sedimentation coefficients and given the prediction of C-terminal coiled-coils for SEPT2, SEPT6 and SEPT7 (de Almeida Marques et al., 2012; Low and Macara, 2006; Sala et al., 2016) (Fig. 1 A, B), we used coiled-coil modeling and homology-modeling software to build a model of the SEPT2-SEPT6-SEPT7-SEPT9-SEPT9-SEPT7-SEPT6-SEPT2 octamer (Fig. 1 H). We used this model structure together with the Svedberg equation and the HullRad algorithm (Fleming and Fleming, 2018) which calculates hydrodynamic properties of molecules from their structures, to obtain the theoretical diffusion coefficients of octamers and tetramers, as well as their theoretical sedimentation coefficients (table in Fig. 1 G, see Materials and methods for details). The experimental sedimentation coefficients for octamers-9_i1 and octamers-9_i3 were in excellent agreement with the ones estimated from the model structure, with the sedimentation coefficient distributions again providing no evidence for the presence of tetramers that are expected to sediment much more slowly (by ∼ 3 S).

### Single particle electron microscopy analysis of recombinant septin octamers reveals the flexibility of N- and C-terminal extensions, and provides evidence for SEPT2 and SEPT9 occupying the end and central positions, respectively

To visualize the isolated octamers, we employed single particle electron microscopy (EM) of negative-stained octamer preps in a high salt buffer (300 mM KCl) to prevent septin complexes from polymerizing. Low magnification EM images of negative-stained octamer preps highlighted the rod-like appearance of the complexes (Fig. 2 A). Single particles in such fields, typically ∼3-4,000 particles, were computationally aligned and classified into classes with distinct features (for example, orientation, curvature, or number of subunits). Each class typically contained ∼50-100 particles (see Materials and methods). Fig. 2 B shows a gallery of class averages for octamers-9_i3. Each image is the average of all the particles in a given class, and has an increased signal to noise ratio compared to the raw images, which allows us to distinguish individual septin subunits within the octameric complex. All class averages for octamers-9_i3 exhibited a characteristic rod shape, similarly to recombinant human/mammalian septins (Mavrakis et al., 2016; Mendonca et al., 2019; Sirajuddin et al., 2007) or septins isolated from mammalian cell lines and tissues (Hsu et al., 1998; Kim et al., 2011; Kinoshita et al., 2002; Sellin et al., 2011; Soroor et al., 2020), and in line with rod-shaped septin complexes from other species, including budding yeast (Bertin et al., 2008; Frazier et al., 1998; Garcia et al., 2011; Taveneau et al., 2020), *C. elegans* (John et al., 2007) and *Drosophila* (Akhmetova et al., 2015; Field et al., 1996; Mavrakis et al., 2014). The class averages did not show additional densities at their ends or along their sides, suggesting an intrinsic orientational flexibility in the junction between the G domain and the coiled-coils of SEPT2, SEPT6 and SEPT7, whose densities are averaged out in such analysis. Such flexibility for the coiled-coils, deduced from the absence of electron density in the crystal structure of the SEPT2-SEPT6-SEPT7 trimer (Sirajuddin et al., 2007) and the absence of additional densities in single particle EM of budding yeast, mammalian, *C*.*elegans* and *Drosophila* complexes (Bertin et al., 2008; Garcia et al., 2011; John et al., 2007; Mavrakis et al., 2014; Mavrakis et al., 2016; Mendonca et al., 2019; Taveneau et al., 2020), seems to be a general feature of septins. Moreover, there was no density that could be assigned to the large (∼27-kDa) N-terminus of the long isoforms, in line with the absence of any secondary structure prediction for this SEPT9 long isoform-specific domain (see Materials and methods).

**Figure 2.**
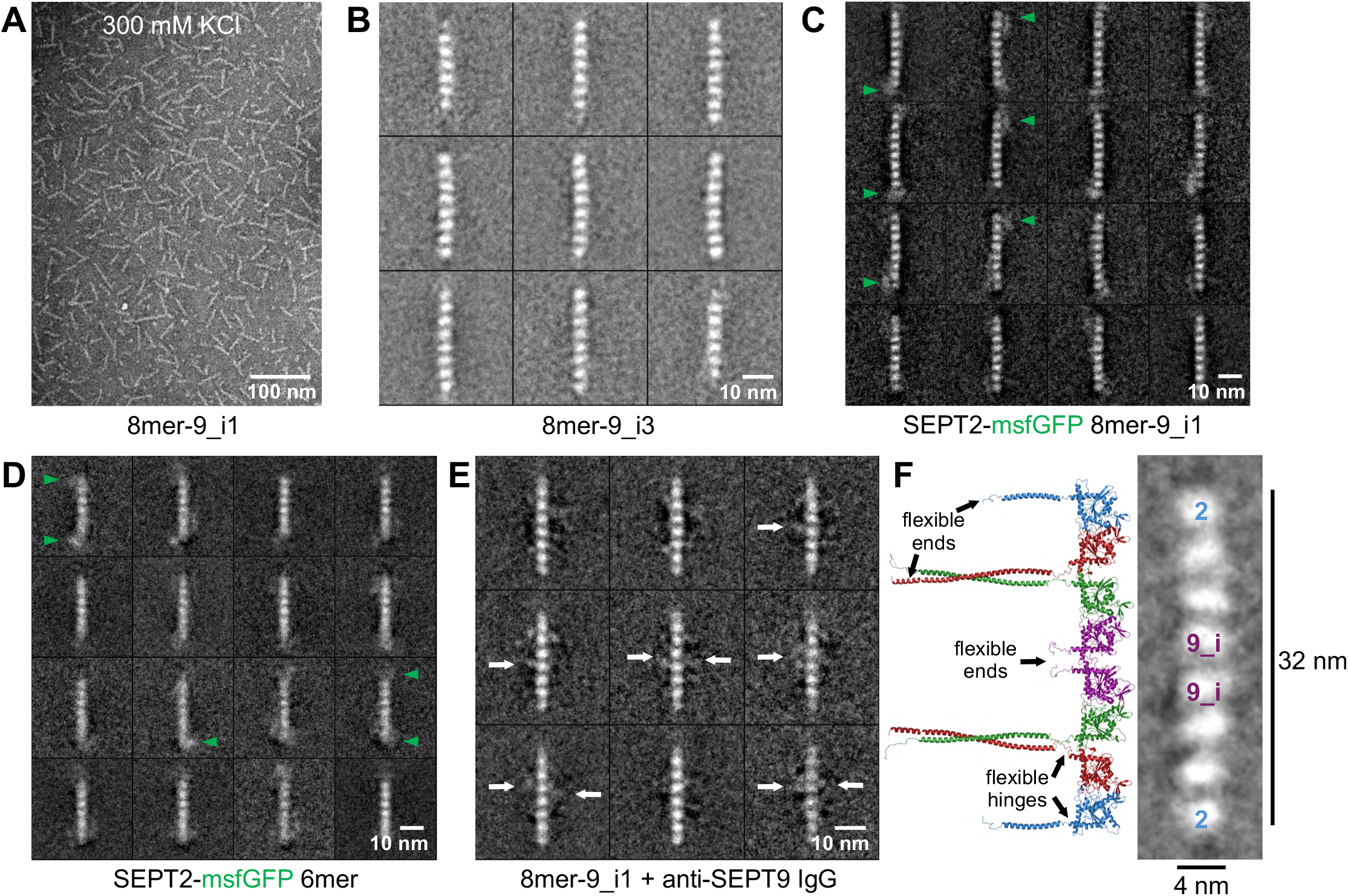
Electron microscopy analysis of recombinant SEPT9-containing octamers. **(A)** Negative-stain EM image of recombinant 8mer-9_i1 at 25 nM in a high salt buffer (300 mM KCl) showing the rod-like appearance of the complexes. **(B-E)** Single particle EM analysis of recombinant septin complexes. Each panel shows a gallery of representative class averages (∼100 particles each) derived from the processing of ∼3,000-4,000 particles from micrographs of negative-stained complexes, as in (A) (see Materials and methods for details). All class averages for 8mer-9_i3 complexes (B) exhibited a rod shape with no evidence of additional densities at their ends or along their sides, consistent with a high degree of flexibility for the coiled-coils of SEPT2, SEPT6 and SEPT7, and in line with the absence of secondary structure prediction for the large (27-kDa) N-terminus of the long SEPT9 isoforms. Class averages of SEPT2-msfGFP 8mer-9_i1 (C) and of SEPT2-msfGFP 6mer (D) displayed additional densities at one or both ends of the rods (green arrowheads), indicating that SEPT2 occupies the termini of 6mer and 8mer. The fuzzy density of the C-terminal GFP and its multiple positions around the end subunit suggest an intrinsic orientational flexibility in the junction between the G domain and the coiled-coils of SEPT2. Class averages of 8mer-9_i1 incubated with anti-SEPT9 antibodies (E) exhibited additional fuzzy densities closest to the center subunits (white arrows), consistent with SEPT9 occupying the central positions in the octameric rods. **(F)** shows the juxtaposition of the model of the octamer (Fig. 1 H) to a high magnification class average image of an octamer from (B). All septin complexes shown contain full-length, human septins apart from (B) which depicts an example of mammalian septin octamers-9i3 containing mouse SEPT2-, human SEPT6-, human SEPT7ΔN19 and human SEPT9_i3 (see Materials and methods).

Class averages of SEPT2-msfGFP octamers-9_i1 (Fig. 2 C) did display additional densities at one or both ends of the rods (green arrowheads), indicating that the GFP-tagged SEPT2 subunits occupy the termini of octamers. The same observation was made with SEPT2-msfGFP-containing hexamers (Fig. 2 D). The fuzzy density of the C-terminal GFP and its multiple positions around SEPT2 again point to a flexible hinge region between the last helix of the G-domain (*α*6 helix, Fig. 1 H) and the coiled-coils of SEPT2 (Fig. 1A, B, H). Class averages of octamers-9_i1 incubated with anti-SEPT9 antibodies exhibited additional fuzzy densities closest to the central subunits (Fig. 2 E, white arrows), consistent with SEPT9 occupying the central positions in the octameric rods. Our observations on the positioning of SEPT2 and SEPT9 in the isolated octamers are thus consistent with recent studies on septin positioning in recombinant human hexamers (Mendonca et al., 2019), recombinant SEPT3-containing octamers (DeRose et al., 2020) and SEPT9-containing octamers isolated from cell lysates (Soroor et al., 2020) (Fig. 2 F).

### Recombinant septin octamers harboring either SEPT9_i1, SEPT9_i3, or SEPT9_i5 polymerize into higher-order filament assemblies in solution

In addition to SEPT2 at the ends of octamers, whose presence is determinant for octamer polymerization in solution, other structural elements such as the N- or C-terminal extensions could also impact septin filament assembly (Bertin et al., 2010). To test the effect of the SEPT9-specific N-terminal extension on septin polymerization in solution, we compared octamers containing either a long or a short SEPT9 isoform. To this end, octamers-9_i1, -9_i3 and -9_i5 were either dialyzed or diluted into a low-salt buffer (50 mM KCl). The resulting assemblies were observed with spinning disk fluorescence microscopy on PLL-PEG passivated glass using SEPT2-msfGFP containing octamers (Fig. 3 A-C), and examined at higher spatial resolution by negative-stain EM using nonfluorescent or SEPT2-msfGFP containing octamers (Fig. 3 D-E). Both octamers with a long or a short SEPT9 isoform polymerized into higher-order filament assemblies. Fluorescence microscopy revealed a variety of assembly morphologies: optical sectioning showed that all octamers assembled into interconnected and/or branched networks of straight and curved filament bundles extending along 10-50 μm in the xy plane and by 10-50 μm in z (Video 1 shows a representative z-stack, and Fig. 3 A-C show maximum-intensity projections of z-stacks). Octamers could organize into straight bundles (left panel in Fig. 3 B; Fig. 3 C), but also into what looked like highly convoluted filamentous assemblies (Fig. 3 A; right panel in Fig. 3 B). Measurements of isolated septin filament bundles in solution for octamers-9_i3 and -9_i5 (left panel in Fig. 3 B; Fig. 3 C) showed that they could reach up to ∼5-8 μm in length.

**Figure 3.**
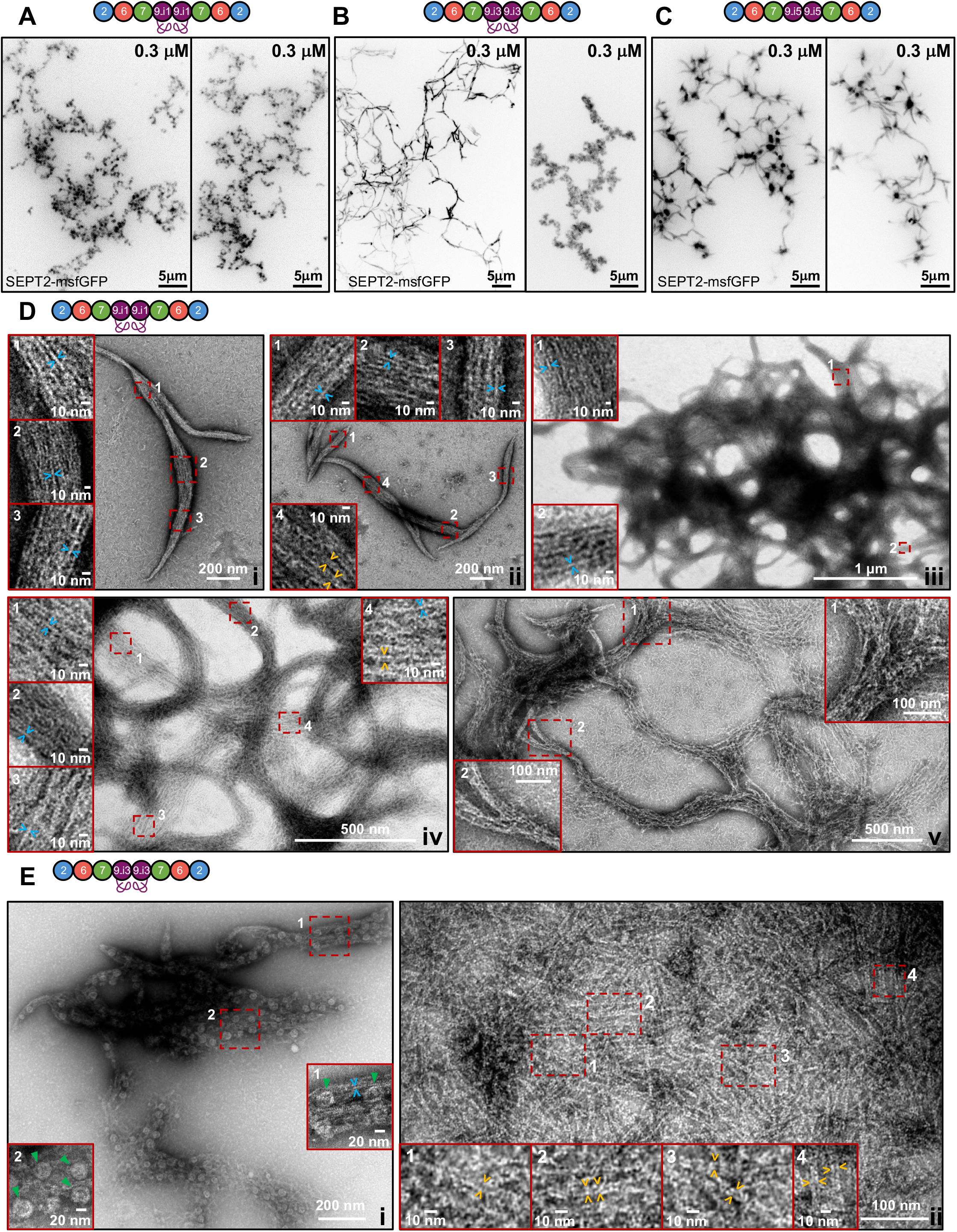
*In vitro* reconstitution of septin polymerization in solution using recombinant septin octamers with distinct SEPT9 isoforms. **(A-C)** Representative spinning disk fluorescence images of higher-order filament assemblies upon polymerization of 8mer-9_i1 (A), 8mer-9_i3 (B) and 8mer-9_i5 (C) after dilution into low salt conditions (50 mM KCl) at the indicated final octamer concentration. Two examples are shown for each. All images shown are maximum-intensity projections and use an inverted grayscale. **(D)** Negative-stain EM images of higher-order filament assemblies upon polymerization of 8mer-9_i1 at low salt (50 mM KCl). 8mer-9_i1 were polymerized at 1 μM (D, i, ii) or 0.2 μM (D, iii-v) final octamer concentration. The insets show magnifications of selected regions of interest (dashed rectangles in red), and highlight single septin filaments (blue arrowheads), possibly paired septin filaments (orange arrowheads), and splayed filament bundles (v). **(E)** Negative-stain EM images of higher-order filament assemblies upon polymerization of 8mer-9_i3 at low salt (50 mM KCl). 8mer-9_i3 were polymerized at 0.2 μM final octamer concentration. The insets show magnifications of selected regions of interest (dashed rectangles in red), and highlight single septin filaments (blue arrowheads), paired septin filaments (orange arrowheads), and wheel-like structures whose perimeter corresponds to two or three octamers connected end to end (green arrowheads). All septin complexes shown contain full-length, human septins apart from the left panel in (E) which depicts an example of mammalian septin octamers-9i3 containing mouse SEPT2-, human SEPT6-, human SEPT7ΔN19 and human SEPT9_i3 (see Materials and methods).

Negative-stain EM similarly revealed a variety of filamentous assemblies. Octamers-9_i1 could organize into isolated or interconnected straight or slightly curved bundles (Fig. 3 D, i-ii), and into networks of interconnected highly convoluted and ring-forming filament bundles (Fig. 3 D, iii-v) corresponding to the similarly convoluted filamentous assemblies in fluorescence microscopy (Fig. 3 A). Septin filament bundles were a few μm long and ∼50-150 nm in width (Fig. S2 E). High magnifications of regions within the filament bundles (red-outlined insets in Fig. 3 D) revealed that bundles were made of single septin filaments (blue arrowheads in Fig. 3 D and measurements of septin filament width in Fig. S2 E) running parallel to each other, with septin filaments within bundles occasionally looking paired (orange arrowheads in Fig. 3 D). Given the high density of filaments and the 2D projection character of negative-stain EM, we cannot conclude if these are truly paired filaments like budding yeast septin filament pairs (Bertin et al., 2008); if so, these would have to be more tightly paired given their interfilament spacing (∼2-3 nm). Septin filament bundles exhibited a high degree of interconnectivity, with a given bundle often showing splayed ends that could connect to one or more different bundles, or with septin filaments forming meshes (Fig. 3 D, v). We speculate that the exposed, flexible coiled-coil-containing C-terminal extensions, and potentially the long N-terminal extensions of SEPT9, drive the interconnections in the higher-order assemblies we observe. Octamers-9_i3 also formed filament bundles (Fig. 3 E, i) and *bona fide* paired filaments with narrower interfilament spacing (∼2-3 nm) than the one observed for budding yeast septin filament pairs in solution (∼10 nm) (Bertin et al., 2008) (Fig. 3 E, ii). Octamers-9_i3 additionally formed wheel-like structures associating with the bundles (Fig. 3 E, i, green arrowheads). These wheels had a diameter of 20-30 nm and could correspond to two or three octamer rods connected end to end; the interior of the wheels appeared to contain electron density. Very similar-looking wheels of similar dimensions have been reported for budding yeast Shs1-containing octamers, with the electron density in their interior attributed to the C-terminal coiled-coils stabilizing these structures (Garcia et al., 2011; Taveneau et al., 2020). Our combined observations from fluorescence and electron microscopy show that recombinant octamers with either a long or a short SEPT9 isoform form higher-order filament assemblies, and are consistent with studies of SEPT9-containing octamers isolated from cell lysates (Soroor et al., 2020).

For comparison, we examined human septin hexamers, mammalian septin hexamers (containing mouse SEPT2, human SEPT6 and human SEPT7ΔN19), and *Drosophila* hexamers whose polymerization was characterized previously (Mavrakis et al., 2014; Mavrakis et al., 2016). Fluorescence microscopy and negative-stain EM of human, mammalian and *Drosophila* hexamers in a low-salt buffer (Fig. S2 A-D) showed that human and mammalian hexamers organized in a very similar manner into straight and curved filament bundles made of single and possibly paired septin filaments (Fig. S2 C, D; Fig. S2 E). *Drosophila* hexamers organized in characteristic needle-like bundles, as reported previously (Mavrakis et al., 2014; Mavrakis et al., 2016), which were not as heavily interconnected as their human counterparts. Human hexamer- and octamer-bundles displayed similar lengths and widths (Fig. S2 E, see legend for median values). The observation of such a large variety in the morphologies of filament assemblies in solution could result from several factors. In this study we examined septin assembly using recombinant octamers bound to the endogenous GDP/GTP in the bacterial cytoplasm, without the exogenous addition of nucleotide during cell lysis or post-purification; it is conceivable that regulation of the GDP/GTP state of septins in cells regulates their higher-order assembly (Weems and McMurray, 2017). A further important element is that interactions of septins with membranes, accessory proteins, or the actin/microtubule cytoskeleton could influence their assembly in cells. Our observations lead us to speculate that, in the absence of any other interacting surface or protein, higher-order septin assembly is dominated by the exposed, flexible coiled-coil-containing C-terminal extensions, resulting in the high plasticity and polymorphism we observe. The filamentous assemblies we observe in solution could reflect higher-order septin filamentous assemblies that have been observed in the cytosol of cells upon perturbation of interacting partners, for example, upon disruption of actin stress fibres (Joo et al., 2007; Kim et al., 2011; Kinoshita et al., 2002; Kinoshita et al., 1997; Xie et al., 1999).

### Recombinant octamers-9_i1, octamers-9_i3 and octamers-9_i5 bind and cross-link actin filaments in solution

Septins are thought to interact with actin filaments either indirectly via myosin-II (Joo et al., 2007), or directly. The possibility of direct interactions between septin hexamers and actin filaments was raised by reconstitution assays showing that recombinant mammalian and *Drosophila* hexamers bind and cross-link actin filaments into bundles (Mavrakis et al., 2014). Given that in cells both long isoforms, SEPT9_i1 and SEPT9_i3, and the short isoform SEPT9_i5 associate with the actin cytoskeleton (Connolly et al., 2011; Dolat et al., 2014b; Kim et al., 2011; Surka et al., 2002; Verdier-Pinard et al., 2017), we sought to test if recombinant octamers-9_i1, -9_i3 and -9_i5 have the capacity to directly bind and cross-link actin filaments, as hexamers do, and if so, whether the presence of a specific SEPT9 isoform makes any difference.

To this end, we used spinning disk fluorescence microscopy to image dilute solutions (1 μM) of single actin filaments on PLL-PEG passivated glass, after spontaneous polymerization of purified rabbit muscle G-actin in the presence or absence of nonfluorescent or SEPT2-msfGFP-containing octamers-9_i1, -9_i3 and -9_i5 (Fig. 4 A-E). In the absence of octamers, fluorescence microscopy showed isolated fluctuating single actin filaments, as expected (Fig. 4 A and Video 2). In the presence of 0.3 μM SEPT2-msfGFP-containing octamers-9_i1 (Fig. 4 C), octamers-9_i3 (Fig. 4 D), or octamers-9_i5 (Fig. 4 E), fluorescence microscopy revealed actin filaments cross-linked into straight and curved bundles, in a very similar manner for all three types of octamers, and very similarly to cross-linking induced by human, mammalian and *Drosophila* hexamers (Fig. S3 A-C and (Mavrakis et al., 2014)). Actin filament cross-linking was observed for septin concentrations of 20-30 nM and above, with identical results obtained with nonfluorescent septins (data not shown). The images shown in Fig. 4 C-E were captured typically a few hours into actin polymerization or after overnight incubation, and show the coexistence of straight and curved actin filament bundles, either isolated ones or bundles connected to each other forming networks. Thicker actin bundles, corresponding to a brighter signal in the actin channel, fluctuated very little, suggesting that they were rigid, whereas thinner actin bundles and single actin filaments, emanating from the ends or the sides of bundles, or connecting neighboring bundles, were freely fluctuating (Video 3). Septins systematically colocalized with the actin bundles, indicative of their actin filament cross-linking activity (right panels in Fig. 4 C-E depict insets of selected red-outlined regions on the left). Saturating the actin and septin channels to bring out features with weaker signals, notably single actin filaments (arrowheads in the actin channel in Fig. 4 C-E), revealed that septins localized only to actin bundles and not to single actin filaments, suggesting cooperativity in septin-actin binding. Such cooperativity has also been reported for other actin filament cross-linkers (Winkelman et al., 2016). The similarities in the actin filament cross-linking capacities of human octamers-9_i1, -9_i3 and -9_i5 and human hexamers raise the possibility that SEPT9, in the context of an octameric complex, does not contribute to actin filament cross-linking, but we cannot exclude such a contribution (Dolat et al., 2014b; Smith et al., 2015). In the latter case, the contribution of SEPT9 to actin filament cross-linking seems to be indistinguishable from the one of the other septins in the complex. Our observations raise the possibility that septin-decorated actin structures such as stress fibres and the cytokinetic ring reflect direct interactions between actin filaments and hexameric or/and octameric septin complexes.

**Figure 4.**
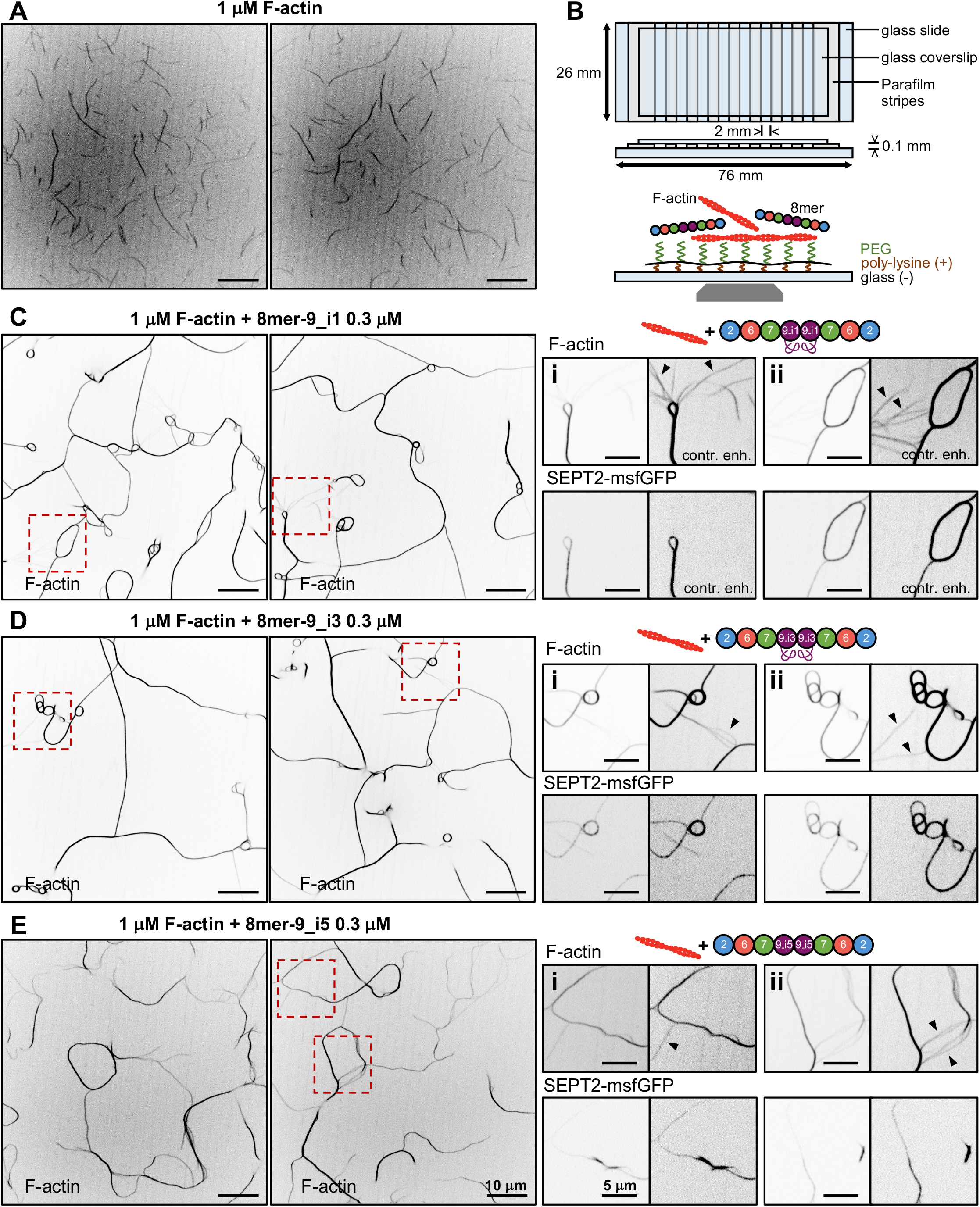
*In vitro* reconstitution of actin filament cross-linking by recombinant human septin octamers with distinct SEPT9 isoforms. **(A-B)** Representative spinning disk fluorescence images of reconstituted, fluctuating single actin filaments (A) upon spontaneous polymerization of G-actin at 1 μM on PLL-PEG-passivated glass in a flow chamber constructed as shown in (B) (see Materials and methods for details). Actin filaments are visualized by including AlexaFluor568-conjugated phalloidin during polymerization. The time lapse sequence containing the still image shown on the left panel of (A) is shown in Video 2. **(C-E)** Representative spinning disk fluorescence images of reconstituted actin filaments, prepared as in (A), and polymerized in the presence of 8mer-9_i1 (C), 8mer-9_i3 (D), or 8mer-9_i5 (E). Actin filaments are visualized with AlexaFluor568-conjugated phalloidin, and septins with SEPT2-msfGFP. Two examples of large fields of view are shown for each, depicting the similar cross-linking of actin filaments into actin filament bundles in the presence of all three types of octamers; only actin labeling is shown. A time lapse sequence containing the still image shown on the right panel of (C) is shown in Video 3. Insets on the right side of each panel show higher magnifications of selected regions of interest on the left (dashed squares in red). Two regions of interest (i, ii) are shown in each case, depicting both the actin (top row) and septin (bottom row) signals. For each inset, actin and septin signals are shown in duplicates: the first set shows the raw signals without any saturation, whereas the second set, adjacent to the first one, shows both actin and septin signals after deliberate contrast enhancement. The contrast-enhanced images in the actin channel saturate the actin bundles, while bringing out weaker-intensity single actin filaments (black arrowheads). The respective contrast-enhanced images in the septin channel show the presence of septins in actin bundles, but their absence from single actin filaments. Scale bars in all large fields of views, 10 μm. Scale bars in all insets, 5 μm. All images shown use an inverted grayscale.

In summary, our study describes the first isolation and characterization of recombinant human SEPT2-SEPT6-SEPT7-SEPT9-SEPT9-SEPT7-SEPT6-SEPT2 octamers containing distinct long or short SEPT9 isoforms. The employed two-tag purification scheme is rapid, taking one full day starting from bacteria lysis, works with prepacked columns, batch resins, or combinations thereof thus providing flexibility, and yields ∼1-3 mg of purified octamers (a few hundreds of microliters in the micromolar concentration range), enabling biochemical and biophysical reconstitution studies at physiological septin concentrations. A combination of biochemical and biophysical assays confirmed the octameric nature of the isolated complexes in solution, and also provided evidence for SEPT2 and SEPT9 occupying the end and central positions in the octamer. Recombinant octamers with either a long or a short SEPT9 isoform were competent for polymerization in solution, and they all shared the capacity to cross-link actin filaments. Given the importance of SEPT9 in mammalian physiology and disease, the isolation of recombinant human septin octamers bearing distinct SEPT9 isoforms will facilitate studies of SEPT9 in the physiological context of its assembly into hetero-octamers. Septins engage in multiple interactions, making it difficult to dissect their function in the complex environment of the cell. Cell-free reconstitution studies with SEPT9-containing octamers and candidate interacting partners thus provide a powerful complementary approach to cellular and animal model studies for exploring human septin function.

## Materials and methods

### Plasmids and cloning

We refer to the mammalian septin protein products as SEPT2, SEPT6, SEPT7, and SEPT9 (SEPT9_i to denote isoforms; the longest isoform being i1, the next longest i2, and so on), nonitalicized and with all letters capitalized, according to the mammalian septin nomenclature established by (Hall et al., 2008; Macara et al., 2002). Human septin gene symbols are italicized with all capital letters, and mouse septin gene symbols italicized with their first letter capitalized. Mouse SEPT2, human SEPT6 and human SEPT7ΔN19 cDNAs were originally obtained from A. Wittinghofer (Max Planck Institute of Molecular Physiology, Germany) and used for the expression and purification of recombinant mammalian SEPT2-, SEPT6-, SEPT7-containing hexamers bearing TEV-cleavable His_6_-, N-terminally-tagged mouse SEPT2, human SEPT6 and noncleavable Strep-tag-II-, C-terminally-tagged human SEPT7ΔN19 (Fig. S2 D and Fig. S3 C) (Mavrakis et al., 2014). Human SEPT9_i1 cDNA was a gift from C. Montagna (Albert Einstein College of Medicine, USA). Human SEPT9_i3 cDNA was a gift from W. Trimble (University of Toronto, Canada). Both human SEPT9_i1 and SEPT9_i3cDNAs have a valine at position 576 for SEPT9_i1 and 558 for SEPT9_i3 instead of a methionine, compared to the respective sequences in UNIPROTKB (identifiers Q9UHD8-1 and Q9UHD8-2); a valine in that position is found in several human clones and many other primates suggesting this is a polymorphism. We used the pET-MCN vectors pnEA-vH (pET15b backbone) and pnCS (pCDF-DuET backbone) for subcloning (Diebold et al., 2011).

Building on our cloning strategy for generating hexamers and with the plasmids available at the time (Mavrakis et al., 2014; Mavrakis et al., 2016), we generated initially a bicistronic pnEA-vH vector for co-expression of TEV-cleavable His_6_-, N-terminally-tagged mouse SEPT2 and human SEPT6, and a bicistronic pnCS vector for co-expression of human SEPT7ΔN19 and noncleavable Strep-tag-II-, C-terminally-tagged human SEPT9_i3. To this end, we digested human SEPT6 in pnCS with SpeI/XbaI, ligated the insert to SpeI-digested mouse SEPT2 in pnEA-vH and selected the clones with the correct insert orientation using restriction analysis. Similarly, we digested human SEPT9_i3 in pnCS with SpeI/XbaI, ligated the insert to SpeI-digested human SEPT7ΔN19 in pnCS and selected the clones with the correct insert orientation using restriction analysis. The combination of these vectors was used to produce and purify recombinant mammalian SEPT9_i3-containing octamers bearing TEV-cleavable His_6_-, N-terminally-tagged mouse SEPT2, human SEPT6, human SEPT7ΔN19 and noncleavable Strep-tag-II-, C-terminally-tagged human SEPT9_i3 (Fig. 2 B; left panel in Fig. 3 E).

To introduce a TEV cleavage site for Strep-tagged SEPT9_i3 and also generate SEPT9_i1 and SEPT9_i5-containing octamers, and to introduce the missing N-terminal 19 residues in SEPT7 (Fig. S1 B), SEPT7ΔN19 being initially erroneously annotated as full-length (Macara et al., 2002), we proceeded as follows. To generate full-length human SEPT7 (UNIPROTKB identifier Q1681-1), we linearized the pnCS plasmid with NdeI/NheI and employed seamless cloning (In-Fusion HD Cloning Plus Kit from Takara Bio, Cat. # 638910) with the following primers: forward 5’-AAGGAGATATACATATGTCGGTCAGTGCGAGATCCGCTGCTGCTGAGGAGAGGAGCGTCAACAGCAGCACCATGGTAGCTCAACAGAAGAACCTTG-3’ and reverse 5’-GCAGCCTAGGGCTAGCTCTAGACTATTAGGATCcTTAAAAGATCTTCCCTTTCTTCTTGTTCTTTTCC-3’. To insert human SEPT9_i1, SEPT9_i3, and SEPT9_i5 with a TEV-cleavable C-terminal Strep-tag-II upstream of SEPT7, we linearized the SEPT7-containing plasmid with SpeI and used seamless cloning with the following primers: forward 5’-ACAATTCCCCACTAGTAATAATTTTGTTTAACTTTAAGAAGGAGATATACATATGAAGAAGTCT-3’ for SEPT9_i1, forward 5’-ACAATTCCCCACTAGTAATAATTTTGTTTAACTTTAAGAAGGAGATATACAtATGGAG AGG-3’ for SEPT9_i3 and forward 5’-ACAATTCCCCACTAGTAATAATTTTGTTTAACTTTAAGAAGGAGATATACATATGGCCGACACCCCCAG-3’ for SEPT9_i5 and reverse 5’-CAAAATTATTACTAGTTTATTTTTCGAACTGCGGGTGGCTCCAGCCGCTGCCGCTG CCCTGGAAGTAAAGGTTTTCCATCTCTGGGGCTTCTGGC-3’ thus generating bicistronic pnCS vectors for co-expression of full-length human SEPT7 and human SEPT9_i (Fig. S1 A, B).

Mouse and human SEPT2 differ in 5 residues (I67V, S207N, S352G, S354G, Q359H, the latter residue being human), presenting 98.61% identity. To replace mouse SEPT2 with human SEPT2 in the bicistronic pnEA-vH vector for co-expression of TEV-cleavable His_6_-, N-terminally-tagged mouse SEPT2 and human SEPT6 (Fig. S1 A, B), we linearized pnEA-vH with KpnI/NheI and employed seamless cloning with the following primers: forward 5’-ATCATCACAGCAGCGGTACCGGCAGCGGCGAAAACCTTTACTTCCAGGGCCATATGTCTAAGCAACAACCAACTCAGTTTATAAATC-3’ and reverse 5’-ATCTCCTAGGGCTAGCTCTAGACTATTAGGATCCTCACACATGGTGGCCGAGAG-3’. The human SEPT2 cDNA containing several restriction sites routinely used in cloning (KpnI, NheI, BamHI) and to facilitate subsequent subcloning using these sites, we chose to generate a synthetic human SEPT2 coding sequence (Eurofins Genomics, Germany) that employs the codon usage of mouse SEPT2 (which does not contain the mentioned restriction sites) except for the five codons that differ between the two species, for which we used codons encoding the human residues V67, N207, G352, G354, H359.

To produce fluorescent octamers, we swapped SEPT2 in the dual expression vector for SEPT2 with its C-terminus tagged with monomeric (V206K) superfolder GFP (msfGFP) (no linker sequence). We generated a synthetic msfGFP coding sequence (Eurofins Genomics, Germany), linearized the dual expression pnEA-vH plasmid with KpnI/NheI and employed two-insert seamless cloning with the following primers: forward 5’-ATCATCACAGCAGCGGTACCGGCAGCGGCGAAAACCTTTACTTCCAGGGCCATATGTCTAAGCAACAACCAACTCAGTTTATAAATC-3’ and reverse 5’-TTGGACACCACATGGTGGCCGAGAGC-3’ for SEPT2, and forward 5’-CCATGTGGTGTCCAAGGGCGAGGAGC-3’ and reverse 5’-ATCTCCTAGGGCTAGCTCTAGACTATTAGGATCCTTACTTGTACAGCTCATCCATGC CCAG-3’ for msfGFP.

To generate recombinant human hexamers bearing TEV-cleavable His_6_-, N-terminally-tagged human SEPT2, human SEPT6 and TEV-cleavable Strep-tag-II-, C-terminally-tagged human SEPT7 (Fig. S1 A, B; Fig. S2 A, C; Fig. S3 A) we employed a similar cloning strategy. To insert human SEPT2 or human SEPT2-msfGFP in the pnEA-vH vector, we linearized pnEA-vH with KpnI/NheI and employed seamless cloning with the same primers used above in the context of the dual vector. To generate TEV-cleavable Strep-tag-II-, C-terminally-tagged full-length human SEPT7, we linearized the pnCS plasmid with NdeI/NheI and employed seamless cloning with the following primers: forward 5’-AAGGAGATATACATATGTCGGTCAGTGCGAGATCCGCTGCTGCTGAGGAGAGGAGCGTCAACAGCAGCACCATGGTAGCTCAACAGAAGAACCTTG-3’ and reverse 5’-GCAGCCTAGGGCTAGCTCTAGACTATTAGGATCCTTATTTTTCGAACTGCGGGTGG CTCCAGCCGCTGCCGCTGCCCTGGAAGTAAAGGTTTTCAAAGATCTTCCCTTTCTTCTTGTTCTTTTCC-3’. To insert human SEPT6 upstream of SEPT7, we linearized the SEPT7-containing plasmid with SpeI and used seamless cloning with the following primers: forward 5’-ACAATTCCCCACTAGTAATAATTTTGTTTAACTTTAAGAAGGAGATATACATATGGCAGCGACCGATATAGC-3’ and reverse 5’-CAAAATTATTACTAGTCTATTAGGATCCTTAATTTTTCTTCTCTTTGTCTCTCTTCAGAGTCTGTGAG-3’ thus generating a bicistronic pnCS vectors for co-expression of human SEPT6 and full-length human SEPT7.

Recombinant *Drosophila* hexamers bearing TEV-cleavable His_6_-, N-terminally-tagged DSep1, untagged or mEGFP-, N-terminally-tagged DSep2 and noncleavable Strep-tag-II-, C-terminally-tagged Peanut were described previously (Fig. S2 B) (Mavrakis et al., 2014; Mavrakis et al., 2016). To generate *Drosophila* hexamers along the lines of the human ones, i.e. with a TEV-cleavable Strep-tag for Peanut and with the C-terminus of DSep1 (the human SEPT2 homolog) tagged with msfGFP (Fig. S2 B; Fig. S3 B), we proceeded as follows. To insert DSep1-msfGFP in the pnEA-vH vector, we linearized pnEA-vH with NdeI/BamHI and employed two-insert seamless cloning with the following primers: forward 5’-ACTTCCAGGGCCATATGGCCGATACAAAGGGCTTTTC-3’ and reverse 5’-TTGGACACCTGCTGGGCCTGCATGC-3’ for DSep1, and forward 5’-CCAGCAGGTGTCCAAGGGCGAGGAGC-3’ and reverse 5’-TAGACTATTAGGATCCTTACTTGTACAGCTCATCCATGCCCAG-3’ for msfGFP. To introduce the TEV cleavage site for Peanut in the bicistronic pnCS vector for co-expression with DSep2, we linearized the dual expression pnCS plasmid with NcoI/NheI and employed seamless cloning with the following primers: forward 5’-CGCCAGAAGCCCATGGAG-3’ and reverse 5’-GCAGCCTAGGGCTAGCTCTAGACTATTAGGATCCTTATTTTTCGAACTGCGGGTGGCTCCAGCCGCTGCCGCTGCCCTGGAAGTAAAGGTTTTCGAACAGACCCTTCTTTTTCTTCTCCTTCTTGC-3’.

All primers for seamless cloning were Cloning Oligo (<60 bp) or EXTREmer (>60 bp) synthesis and purification quality from Eurofins Genomics, Germany. All restriction enzymes were FastDigest enzymes from Thermo Scientific. All plasmids were verified by sequencing (Eurofins Genomics, Germany) after each cloning step, including the midipreps used for protein production.

### Production and purification of recombinant human and *Drosophila* septin complexes

pnEA-vH and pnCS plasmids were co-transformed into *E. coli* BL21(DE3), and co-transformants selected on LB agar plates with carbenicillin and spectinomycin each at 100µg/mL. A single colony was selected to prepare an overnight preculture at 37°C with LB medium containing antibiotics at 100µg/mL; the volume of the preculture was 1/50 of the final culture volume. Terrific broth with antibiotics at 50µg/mL was inoculated with the pre-culture and incubated at 37°C. We typically prepared 3.5-5 L of culture: the culture volume per Erlenmeyer flask was 1/3 of the flask volume to allow for efficient oxygenation. For nonfluorescent septins we let bacteria grow to A_600nm_ ∼ 2 before inducing expression with 0.5 mM IPTG for 3 h at 37°C. For fluorescent septins we let bacteria grow to A_600nm_∼ 0.6-0.8 before inducing expression with 0.5 mM IPTG for overnight expression at 17°C; the incubator was protected from light with aluminum foil in this case. The culture was stopped by centrifuging at 3,400 g for 15 min and 4°C, and the supernatant used to pool all bacteria pellets in 50-mL Falcon tubes, which were further centrifuged at 5,000 g for 10 min and 4°C. Bacteria pellets were stored at −20°C until protein purification. Bacteria expressing GFP-tagged septins yield yellow-greenish pellets.

On the day of purification we resuspended the pellet in ice-cold lysis buffer (50 mM Tris-HCl pH 8, 300 mM KCl, 5 mM MgCl_2_, 0.25 mg/mL lysozyme, 1 mM PMSF, cOmplete™ protease inhibitor cocktail (1 tablet per 50 mL), 10 mg/L DNase I, 20 mM MgSO_4_) using gentle agitation for 30 min at 4°C, and lysed cells on ice using a tip sonicator with 5 cycles of 30 s “ON”, 15 s “OFF”. We typically use 100 mL of lysis buffer for a starting 3.5-5 L culture. The lysate was clarified by centrifugation for 30 min at 20,000 g and 4°C, the supernatant loaded on a StrepTrap HP column equilibrated with 50 mM Tris-HCl pH 8, 300 mM KCl, 5 mM MgCl_2_ and the column washed with the same buffer. Strep-tag-II-containing septin complexes were eluted with 50 mM Tris-HCl pH 8, 300 mM KCl, 5 mM MgCl_2_, and 2.5 mM desthiobiotin: desthiobiotin was prepared fresh right before use. All fractions contained in the elution peak were collected. The pooled fractions were then loaded to a HisTrap HP column equilibrated with 50 mM Tris-HCl pH 8, 300 mM KCl, 5 mM MgCl_2_, the column washed with the same buffer, and His_6_-tag-containing complexes eluted with 50 mM Tris-HCl at pH 8, 300 mM KCl, 5 mM MgCl_2_, and 250 mM imidazole. We typically do not concentrate the protein further, thus we collect only the highest-concentration peak fractions (∼0.6 − 1.2 mL). Both affinity column steps were performed on an ÄKTA pure protein purification system (Cytiva). To remove imidazole we either performed an overnight dialysis step or used a PD-10 column, also including DTT in this last step. The final elution buffer, in which septins are stored, was 50 mM Tris-HCl pH 8, 300 mM KCl, 5 mM MgCl_2_, and 1 mM DTT. All purification steps were performed at 4°C. We typically purify septin complexes in a single day (starting from cell lysis) to minimize unnecessary exposure to proteases and contaminants and maintain protein integrity and functionality.

Protein concentrations were assessed with absorbance measurements at 280 nm using the calculated extinction coefficients for the respective complexes, and 20- or 50-μL aliquots were flash-frozen in liquid nitrogen and stored at −80°C until further use. This protocol yielded typically 0.5-1 mL of purified octamers at ∼1-3 mg/mL (∼2-6 μM) and 0.5-1 mL of purified hexamers at ∼1-3 mg/mL (∼3-9 μM) from a starting 3.5 L of bacterial culture. Extinction coefficients and molecular masses used for concentration conversions were computed from the primary amino acid sequences using ExPASy (http://web.expasy.org/protparam/) and considering two copies of each full-length septin, tags included, and are summarized in the following table. The calculation of these parameters for mammalian and *Drosophila* hexamers were described in (Mavrakis et al., 2016).

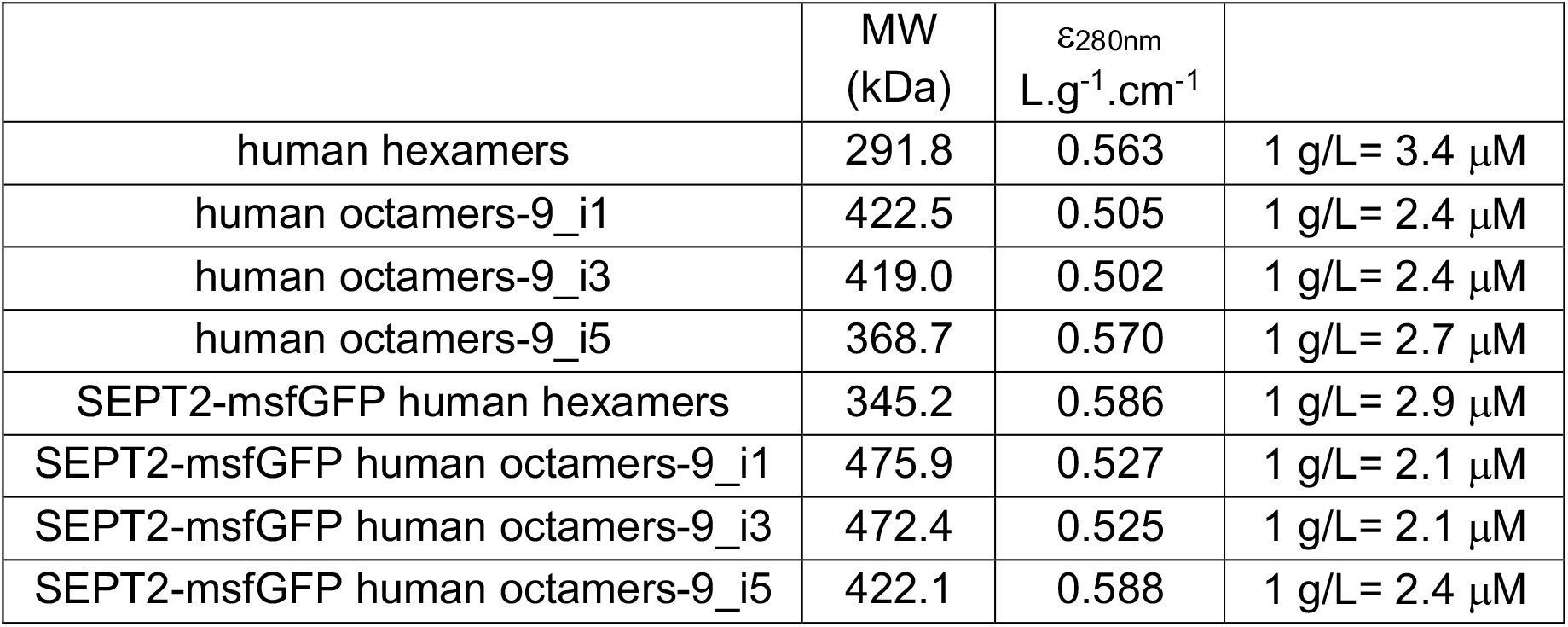

Septin preps were used closest to their purification and typically within 2-3 months upon storage at −80°C. Septin complexes were typically used within 3-4 days upon thawing and not frozen back. Electron microscopy of the purified septin complexes did not show significant aggregation necessitating a gel filtration step, thus size exclusion chromatography used previously (Mavrakis et al., 2014; Mavrakis et al., 2016) was not included. Purified protein was examined as such for EM, whereas it was systematically airfuged right before use for reconstitution assays using fluorescence microscopy to pellet any remaining or formed aggregates upon freezing (see respective methods below). This purification protocol was validated and is routinely used in three different labs (I. Fresnel, I. Curie, TU Delft) with slight variations that do not affect the final result. For example, the order of StrepTrap HP and HisTrap HP columns is inverted, or the nickel-affinity step uses batch resin instead of prepacked columns. This production and purification protocol was used to purify at least six preps of human septin octamers-9_i1, six preps of human septin octamers-9_i3, one prep of human septin octamers-9_i5, and six preps of human septin hexamers. The purification of mammalian septin octamers-9_i3, containing mouse SEPT2, human SEPT6, human SEPT7ΔN19, and human SEPT9_i3 (Fig. 2 B and left panel in Fig. 3 E) was done as described previously (Mavrakis et al., 2014; Mavrakis et al., 2016): the main difference with respect to the protocol described above is that the columns were inverted, nickel affinity used as a first step, and that we employed an additional gel filtration step.

### Materials and reagents for septin complex production and purification

The sources and identifiers for the chemicals used for recombinant protein production and purification are as follows. *E. coli* BL21(DE3) competent cells from Agilent (200131) or Thermo Scientific (EC0114). Carbenicillin (for pnEA-vH selection) from SIGMA (C3416), Condalab (6803), or Fisher Scientific (Fisher Bioreagents BP2648-1). Spectinomycin (for pnCS selection) from Sigma (S4014) or Fisher Scientific (Fisher Bioreagents BP2957-1). LB broth medium from Sigma (L3022) or Condalab (1231). LB agar from Sigma (L2897) or Condalab (1083). SOC medium from Sigma (S1797) or Fisher Scientific (Fisher Bioreagents BP9740). Terrific Broth from MP Biomedicals (091012017) or Fisher Scientific (Fisher Bioreagents BP2468-10). IPTG from Euromedex (EU0008-C). Imidazole with low absorbance at 280 nm from Fisher Scientific (Fisher Chemical I/0010/53). PMSF from Sigma (78830). Lysozyme from Euromedex (5933). cOmplete™ Protease Inhibitor Cocktail Tablets from Sigma (Roche, 11836145001). DNase I from Sigma (Roche, 10104159001). *d*-Desthiobiotin from Sigma (D1411). DTT from Sigma (D0632). HisTrap HP 1 mL columns from Cytiva (17524701). StrepTrap HP 1 mL columns from Cytiva (28907546). 20K MWCO Slide-A-Lyzer cassettes from Thermo Scientific (87735). PD-10 desalting columns from Cytiva (17085101).

### SDS-PAGE and Western blot

We assessed septin prep purity and protein integrity by SDS-PAGE and Western blot. Purified septin complexes were analyzed by SDS-PAGE using 4-20% precast polyacrylamide gels (Mini-PROTEAN TGX Gels from Bio-Rad, 4561095), or hand-casted 12% or 15% Tris-glycine polyacrylamide gels, followed by staining with InstantBlue Coomassie-stain (Expedeon, ISB1L) (Fig. S1 C-E). Molecular weight markers were Precision Plus Protein All Blue Standards from Bio-Rad (1610373) in all gels shown apart from Fig. S1 C (Pierce Unstained Protein MW Marker from Thermo Scientific, 26610) and Fig. S1 D (PageRuler Plus Prestained Protein Ladder from Thermo Scientific, 26619). SDS-PAGE was performed for each septin prep.

Western blots were performed for 2 different preps for each nonfluorescent and SEPT2-msfGFP hexamer and 2 different preps for each nonfluorescent and SEPT2-msfGFP long SEPT9 isoform-containing octamer, with similar results. Gels for Western blot detection were loaded with 10 ng of purified protein. Gel, transfer membrane (Immobilon-P^SQ^ membrane, Sigma ISEQ85R), filter pads and filter papers were incubated in transfer buffer (25 mM Tris, 192 mM glycine, 20% methanol) for 15 minutes before assembly in the Mini Trans-Blot transfer cell (Bio-Rad, 1703935). The transfer was done for 16 h at 4°C and at 110 mA constant current. The membrane was then blocked in a 5% w/v dry nonfat milk TBS-T solution (20 mM Tris-HCl pH 7.5, 200 mM NaCl, 0.1 % v/v Tween20) for 2 h under constant agitation. Primary and secondary antibodies were diluted in the same blocking solution and incubated over the membrane for 60 min each under constant agitation. Between antibody incubations, membranes were washed 3 times for 10 min with TBS-T, the very last wash before detection only with TBS. To detect specific septins in recombinant human hexamers and octamers, we used rabbit anti-SEPT2 (1:2,500, Sigma, HPA018481), rabbit anti-SEPT6 (1:1,500, Santa Cruz Biotechnology, sc-20180), rabbit anti-SEPT7 (1:200, Santa Cruz Biotechnology, sc-20620), rabbit anti-SEPT9 (1:1,500, Proteintech, 10769-1-AP), and HRP-conjugated anti-rabbit antibody (1:10,000, Cytiva, NA934). Chemiluminescent detection was done with an Amersham ImageQuant 800 imager (Cytiva, 29399481) using Amersham ECL Select Western Blotting Detection Reagent (Cytiva, RPN2235) diluted 5 times in Milli-Q water. The membrane was incubated with the diluted reagent for 30 s, and washed for 10 s in TBS right before image acquisition. Images were collected in time series mode every 10 s, for a total of 50 images, and processed with ImageQuantTL software for molecular size calculation. In 4-20% Tris-glycine gels, the apparent mass of SEPT6 was larger than its calculated one by ∼3 kDa, resulting in a band right above the one of SEPT7 that migrated as expected. The TEV-Strep-tag-II of SEPT7 in hexamer preps adds 2.2 kDa to the SEPT7 band which thus migrates much closer to SEPT6, making SEPT6 and SEPT7 bands hard to make out. All SEPT9 isoforms migrated much more slowly than their calculated masses: the apparent masses of the two long isoforms were larger by ∼ 12-13 kDa, the one of the short isoform by ∼ 5 kDa. Western blot analysis of hexamer and octamer preps showed that all septins were intact, with the long N-terminal extension of SEPT9 being the most sensitive to proteolysis (Fig. S1 F). The purity and protein integrity of septins in preps, as well as the identification of protein bands in gels were corroborated by mass spectrometry analysis (see respective section below).

### Mass spectrometry analysis and data processing

For analysis of septin hexamers and octamers (one prep for each hexamer and for each long SEPT9 isoform-containing octamer), 1 µg of sample was loaded on 4–12% NuPAGE Novex Bis-Tris polyacrylamide gels (Thermo Scientific, NP0322BOX) and ran for 7 min at 80V to stack proteins in a single band. The gel was further stained with Imperial Protein Stain (Thermo Scientific, 24615), destained in water and proteins cut from the gel. Gel pieces (protein stack or cut protein bands) were subjected to in-gel trypsin digestion after cysteine reduction and alkylation (Shevchenko et al., 2006). Peptides were extracted from the gel and dried under vacuum. Samples were reconstituted with 0.1% trifluoroacetic acid in 2% acetonitrile and analyzed by liquid chromatography (LC)-tandem MS (MS/MS) using a Q Exactive Plus Hybrid Quadrupole-Orbitrap (Thermo Scientific, IQLAAEGAAPFALGMBDK) online with a nanoLC UltiMate 3000 chromatography system (Thermo Scientific, ULTIM3000RSLCNANO). 2 microliters corresponding to 10 % of digested protein were injected in duplicate on the system. After pre-concentration and washing of the sample on a Acclaim PepMap 100 C18 column (2 cm × 100 μm i.d., 100 Å pore size, 5 μm particle size, Thermo Scientific 164564-CMD), peptides were separated on a LC EASY-Spray C18 column (50 cm × 75 μm i.d., 100 Å pore size, 2 µm particle size, Thermo Scientific ES803) at a flow rate of 300 nL/min with a two-step linear gradient (2-20% acetonitrile in 0.1 % formic acid for 40 min and 20-40% acetonitrile in 0.1 % formic acid for 10 min). For peptide ionization in the EASY-Spray source, spray voltage was set at 1.9 kV and the capillary temperature at 250°C. All samples were measured in a data-dependent acquisition mode. Each run was preceded by a blank MS run in order to monitor system background. The peptide masses were measured in a survey full scan (scan range 375-1500 m/z, with 70 K FWHM resolution at m/z=400, target AGC value of 3.00×10^6^ and maximum injection time of 100 ms). Following the high-resolution full scan in the Orbitrap, the 10 most intense data-dependent precursor ions were successively fragmented in the HCD cell and measured in Orbitrap (normalized collision energy of 27 %, activation time of 10 ms, target AGC value of 1.00×10^5^, intensity threshold 1.00×10^4^ maximum injection time 100 ms, isolation window 2 m/z, 17.5 K FWHM resolution, scan range 200 to 2000 m/z). Dynamic exclusion was implemented with a repeat count of 1 and exclusion duration of 10 s.

Raw files generated from mass spectrometry analysis were processed with Proteome Discoverer 1.4.1.14 (Thermo Scientific) to search against the proteome reference of the *Escherichia coli* protein database (4,391 entries, extracted from Uniprot in August 2020). The original fasta file was populated with the sequences of the septin constructs contained in the measured preps. Database search with Sequest HT was done using the following settings: a maximum of two missed trypsin cleavages allowed, methionine oxidation as a variable modification and cysteine carbamidomethylation as a fixed modification. A peptide mass tolerance of 10 ppm and a fragment mass tolerance of 0.6 Da were allowed for search analysis. Only peptides with high Sequest scores were selected for protein identification. False discovery rate was set to 1% for protein identification.

To measure the relative protein abundance in septin preps we employed the Top3 quantitation approach based on the correlation between the sum of the three most intense peptide ions of a given protein and its absolute abundance (Silva et al., 2006). We divided the Top3 value of each identified protein in the protein stack by the sum of all Top3 values, generating a relative Top3 abundance measure, which correlates with the mol fraction of the protein. Septins constituted >97% of the total protein content, with the remaining <3% including GTP cyclohydrolase, biotin carboxylase and several chaperones (DnaK, DnaJ, GrpE, 60 kDa chaperonin). The results for molar fractions down to 0.02% are shown in Fig. S1 G for a hexamer and two octamer preps. The obtained mol fractions of septins, compared with the expected ones for hexamers (33%) and octamers (25%), point to the isolation of stoichiometric hexamers and octamers.

Tryptic peptides were used to identify and assign each septin to the detected bands by Coomassie staining, both for nonfluorescent and SEPT2-msfGFP hexamers and octamers, as shown in Fig. 1 D and Fig. S1 C-E, and septin band assignment was in line with the Western blot analysis (Fig. S1 F). Examples of tryptic peptide coverage for individual septins in recombinant hexamer, octamer-9_i1 and octamer-9_i3 preps are shown in Fig. S1 H, with coverages of 82% (SEPT2), 74% (SEPT6) and 70% (SEPT7) for hexamers, 82% (SEPT2), 70% (SEPT6), 69% (SEPT7) and 80% (SEPT9_i1) for octamers-9_i1, and 82% (SEPT2), 85% (SEPT6), 68% (SEPT7) and 84% (SEPT9_i3) for octamers-9_i3. Tryptic peptides were identified throughout the sequence of each septin, including coiled-coils of all septins and the common N-terminal extension of the long SEPT9 isoforms, which together with the apparent band sizes from SDS-PAGE and the Western blot analysis (Fig. 1D; Fig. S1 F) supports that the isolated septin complexes are intact. Coomassie-stained bands other than the annotated ones in our figures were identified as degradation products of septins or/and contaminants already identified in the analysis of the complexes from protein stacks. Degradation band analysis from different preps suggested that the coiled-coils of SEPT2 and SEPT7 and the N-terminal extension of SEPT9 are most sensitive to proteolysis; the sensitivity of the latter to proteolysis was in line with Western blots using antibodies against the C-terminal half of SEPT9 (Fig. S1 F).

### Native PAGE

Native PAGE was performed on 4-16% NativePAGE Novex Bis-Tris polyacrylamide gels (Thermo Scientific, BN1002BOX) following instructions from the manufacturer. Briefly, two μg of recombinant septin complexes, in their elution buffer (50 mM Tris-HCl pH 8, 300 mM KCl, 5 mM MgCl_2_, 1 mM DTT), were diluted with water and 4x native PAGE sample buffer (Thermo Scientific, BN2003) to achieve a total KCl/NaCl concentration of ∼100 mM and 1x native sample buffer, and were loaded in each gel well. Electrophoresis was performed at 150 V constant voltage until the migration front had reached one third of the gel, when dark cathode buffer was replaced with light anode buffer, then electrophoresis was pursued at 150 V until the migration front had reached the bottom of the gel. Gels were destained in 25% methanol and 10% acetic acid to eliminate most of the background (Coomassie stain from running buffer), then washed twice in pure water for 30 min, placed in Imperial Protein Stain (Thermo Scientific, 24615) for one hour and destained in pure water overnight. Molecular weight standards were NativeMark Unstained Protein Standard (Thermo Scientific, LC0725). Native PAGE was performed for 2 independent preps for human hexamers and for each long SEPT9 isoform-containing octamer, with similar results.

Hexamers migrated with an apparent size of ∼310 kDa, in line with the calculated one (292 kDa). Octamers for both long SEPT9 isoform-containing octamers migrated with apparent sizes of ∼600 kDa, thus much more slowly than their theoretical sizes (423 kDa and 419 kDa, respectively), in line with reported gel filtration and density gradient centrifugation experiments showing that the long SEPT9 isoform-specific N-terminal extension confers a significant increase in the hydrodynamic radius slowing down octamer migration in native PAGE (Sellin et al., 2014).

### Modeling of human septin octamers

Models of human septin octamers were generated in order to analyze and interpret the analytical ultracentrifugation sedimentation velocity experiments. A series of structures of the GTP-binding domains (GBDs) of SEPT2, 6, 7 and 9 have been solved by X-ray crystallography (Rosa et al., 2020; Valadares et al., 2017), with some flexible loops partially missing. The homology modelling software, SWISS-MODEL (Waterhouse et al., 2018), was used to complete the existing GBD structures. To search for templates, SWISS-MODEL uses BLAST (Camacho et al., 2009) and HHblits (Steinegger et al., 2019) for related evolutionary structures matching the target sequence within the SWISS-MODEL Template Library (SMTL version 2020-12-09, last included PDB release: 2020-12-04). For each identified template, the quality of the resulting model is predicted from features of the target-template alignment, and the template with the highest quality is selected for model building using ProMod3. Coordinates which are conserved between the target and the template are copied from the template to the model. Insertions and deletions are remodelled using a fragment library. Finally, the geometry of the resulting model is regularized by using a CHARMM22/CMAP protein force field (Mackerell et al., 2004). The global and per-residue model quality is assessed using the QMEAN scoring function (Studer et al., 2020). Due to the lack of structural information for the long SEPT9 isoform N-terminal extension, the absence of structural homologs for this region (using Phyre2 and SWISS-MODEL) and secondary structure predictions of disorder for this region (using Quick2D), this region was not modeled. Similarly, the lack of structural information for the short N-terminal extensions of SEPT2, 6, and 7, including the *α*0 helices, prompted us not to model these regions. GBD models, starting right after the end of the *α*0 helices and until the end of the *α*6 helices, were generated for SEPT2 37-306 (template PDB 6UPA), SEPT6 40-308 (template PDB 6UPR), SEPT7 48-316 (template PDB 6N0B), and SEPT9 295-568 (template PDB 5CY0; numbering based on SEPT9_i1). To account for the contribution of the predicted coiled-coils in the C-terminal extensions of SEPT2, 6 and 7 to the hydrodynamic properties of the complexes and thus their sedimentation behavior, we extended the GBD models to include the C-terminal domain from the end of the *α*6 helix onwards. Delineation of coiled-coil features was based on secondary structure prediction via Quick2D (Zimmermann et al., 2018). This tool integrates secondary structure predictions from different softwares, including coiled-coil prediction via MARCOIL (Delorenzi and Speed, 2002), PCOILS (Gruber et al., 2006) and COILS (Lupas et al., 1991). The consensus sequences assigned by all three coiled-coil prediction algorithms were used for modeling coiled-coil helices with CCFold software (Guzenko and Strelkov, 2018). The resulting coiled-coils encompass residues 310-349, 321-406, and 336-421 for SEPT2, 6, and 7, respectively.

The models built with SWISS-MODEL and CCFold were still missing the connections between the GBDs and coiled-coils for SEPT2, 6, and 7, the ends of the C-terminal domains right after the predicted CCs, and the C-terminal domain of SEPT9 after the *α*6 helix. Phyre2 (Kelley et al., 2015) was used to construct these flexible parts *ab initio* for SEPT2, 7 and 9; and by homology for the very end of SEPT6 as a structural homolog was found by the software. The different models were generated in the context of the full proteins for higher accuracy. The flexible parts linking the GBDs and coiled-coils and the remaining C-terminal features were isolated from the resulting models with PyMOL (open-source software). GBDs, coiled-coils and flexible parts were then combined with PyMOL. The connections between the GBDs and the coiled-coils for SEPT6 and 7 being of different length and to allow for the aligning of coiled-coil helices of SEPT6 and 7, these connections were stretched out so that they cover the same distance between the GBDs and the start of the coiled-coils without any steric clashes. In the case of SEPT2, the 3 residues between the GBD and the coiled-coil were built directly with PyMOL. Tetrameric SEPT2-SEPT6-SEPT7-SEPT9 and octameric SEPT2-SEPT6-SEPT7-SEPT9-SEPT9-SEPT7-SEPT6-SEPT2 complexes were built by fitting the modeled structures to the crystal structure of the SEPT2-SEPT6-SEPT7 trimer (PDB 2QAG) (Sirajuddin et al., 2007). The PDB files of the modeled tetramers and octamers, with or without the C-terminal extensions, were then used in HullRad to determine their diffusion coefficient and estimate their sedimentation coefficient (see section on analytical ultracentrifugation below). Given that the long SEPT9 isoform N-terminal extensions are not contained in the model structures, the calculated diffusion coefficients were the same for all three SEPT9 isoforms (Fig. 1G and Fig. S1 I).

### Analytical ultracentrifugation

A sedimentation velocity experiment was carried out for one prep each of octamers-9_i1 (0.75 mg/mL) and octamers-9_i3 (0.5 mg/mL), at 40,000 rpm and 20°C in a Beckman Optima XL-A analytical ultracentrifuge, using 12 mm double sector centerpieces in an AN-50 Ti rotor (Beckman Coulter). Scans were acquired in continuous mode at 280 nm, in the absorbance range of 0.1 to 1. The partial specific volume of the proteins and the density and viscosity of the buffer were calculated with SEDNTERP (Laue et al., 1992). At 20°C, the calculated partial specific volume for octamers-9_i1 and −9_i3 was 0.735 mL.g^−1^. The density and viscosity of the buffer (50 mM Tris-HCl pH 8, 300 mM KCl, 5 mM MgCl_2_, 1 mM DTT) were 1.014 g.mL^−1^ and 0.0102 poise, respectively. The data recorded from moving boundaries were analyzed in terms of continuous size distribution functions of sedimentation coefficient, *c(s)*, using the program SEDFIT (Schuck and Rossmanith, 2000) and the apparent sedimentation coefficient at 20°C in water (S_20,W_) determined by peak integration.

A short column sedimentation equilibrium experiment was carried out for one prep each of SEPT2-msfGFP hexamers and SEPT2-msfGFP octamers-9_i5, at 11,000 rpm in a Beckman Optima XL-A analytical ultracentrifuge, using 60 µL of protein loading concentrations from 0.5 to 1.5 mg.mL^−1^, in a six-channel epon charcoal-filled centerpiece in an AN-50 Ti rotor (Beckman Coulter). Septins were in 50 mM Tris-HCl pH 8, 300 mM KCl, 5 mM MgCl_2_, and 1 mM TCEP. Scans were acquired at appropriate wavelengths (280 nm and 485 nm) when sedimentation equilibrium was reached at 4°C. Average molecular masses were determined by fitting a sedimentation equilibrium model for a single sedimenting solute to individual data sets with SEDPHAT.

To determine the theoretical sedimentation coefficient, the PDB file of a given model (tetramer, hexamer or octamer, with or without coiled-coils, and with coiled-coils in different orientations) was analyzed using HullRad (Fleming and Fleming, 2018) to determine the translational diffusion coefficient, *D*. The estimated sedimentation coefficient, *s*, was then obtained using the theoretical molecular mass, *M*, for each complex and the Svedberg equation below, with v the partial specific volume of the protein, *ρ* the solvent density, *R* the gas constant and *T* the temperature:

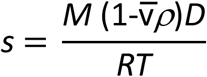

The theoretical sedimentation coefficients calculated in this manner are shown in Fig. 1G and Fig. S1 I. The experimentally measured ones for octamers-9_i1 and -9_i3, and the comparison with the estimated ones in the absence of coiled-coils or in the absence of the long SEPT9 isoform N-terminal extension (SEPT9_i5) are consistent with intact octamers-9_i1 and -9_i3.

### Preparation of flow cells for fluorescence microscopy of *in vitro* reconstituted actin and septins

Microscope glass slides and coverslips were cleaned for 15 min in base-piranha solution (Milli-Q water, 30% ammonium hydroxide, 35% hydrogen peroxide at a 5:1:1 volume ratio), rinsed twice, 5 min each, with Milli-Q water in a bath sonicator, and stored in 0.1 M KOH up to one month. Right before assembling flow cells, slides and coverslips were rinsed twice, 5 min each, with Milli-Q water and dried with synthetic air. Flow cells with ∼10 μL channels were assembled by sandwiching ∼2-mm-wide and ∼2.5-cm-long strips of Parafilm between a cleaned glass slide and coverslip and melting on a hot plate at 120°C (Fig. 4 B). The resulting chambers were passivated by incubating for 45 min with 1 M KOH, rinsing with actin polymerization buffer (see composition in the respective section), incubating for another 45 min with 0.2 mg/mL PLL-PEG, and rinsing with actin polymerization buffer (5 mM Tris-HCl pH 8, 50 mM KCl, 1 mM MgCl_2_, 0.2 mM Na_2_ATP, 1 mM DTT). Flow cells were placed in a Petri-dish along with tissue paper soaked in water to prevent flow channels from drying during the incubation steps and until use.

The sources and identifiers for the materials and chemicals are as follows. Glass slides (26×76 mm) from Thermo Scientific (AA00000102E01FST20). Glass coverslips (24×60 mm) from Thermo Scientific (BB02400600A113FST0). Ammonium hydroxide solution from SIGMA (221228). Hydrogen peroxide solution from SIGMA (95299). PLL-PEG from SuSoS AG (PLL(20)-g[3.5]-PEG(2)).

### Sample preparation for fluorescence microscopy of *in vitro* reconstituted actin and septins

Lyophilized rabbit skeletal muscle G-actin was resuspended to 5 mg/mL (119 μM) in G-buffer (5 mM Tris-HCl pH 8, 0.2 mM Na_2_ATP, 0.1 mM CaCl_2_, 1 mM DTT), aliquots snap-frozen in liquid nitrogen and stored at −80°C. Frozen aliquots were thawed and centrifuged for 30 min at 120,000 g in a benchtop Beckman air-driven ultracentrifuge (Beckman Coulter Airfuge, 340401) to clear the solution from aggregates. Clarified G-actin was kept at 4°C and used within 3-4 weeks.

For reconstitution experiments, G-actin was diluted with G-buffer to 5 μM, and polymerized at 1 μM final concentration in actin polymerization buffer (5 mM Tris-HCl pH 8, 50 mM KCl, 1 mM MgCl_2_, 0.2 mM Na_2_ATP, 1 mM DTT), additionally containing 1 mM Trolox, 2 mM protocatechuic acid (PCA), 0.1 μM protocatechuate 3,4-dioxygenase (PCD) and 0.1% w/v methylcellulose. Trolox and the enzymatic oxygen scavenging system PCA-PCD were used to minimize photobleaching during fluorescence imaging (Cordes et al., 2009; Shi et al., 2010). Methylcellulose was used as a crowding agent to keep actin filaments close to the surface and facilitate their observation. To fluorescently label actin filaments, we polymerized G-actin in the presence of 1 μM Alexa Fluor 568-conjugated phalloidin.

For actin-septin reconstitution experiments, thawed septin aliquots were cleared for 15 min at 120,000 g in a Beckman airfuge right before use. To polymerize G-actin in the presence of septins, we followed the same procedure as above, but mixed G-actin with septins, either nonfluorescent ones or GFP-labeled septins (at 20% GFP molar ratio for octamers-9_i1 and -9_i3, and 100% GFP for octamers-9_i5) to a final septin concentration of 0.3 μM, right before polymerization. Actin and actin-septin samples were prepared with a final volume of 10 μL, were loaded immediately into PLL-PEG-passivated flow channels upon mixing of the components to start polymerization, and flow channels were sealed with VALAP (1:1:1 vasoline:lanoline:paraffin). The contributions of KCl and MgCl_2_ from the septin elution buffer were taken into account to yield the same final composition of actin polymerization buffer. Actin and actin-septin samples were typically incubated overnight at room temperature in the dark before observation. Actin-septin assays were repeated at least four times, using at least two different preps from each nonfluorescent and fluorescent hexamers, 8-mers-9_i1 and 8-mers-9_i3, and one prep from fluorescent 8-mers-9_i5, yielding similar results.

To polymerize septins in the absence of actin, we followed the same procedure as above, but replaced the G-actin solution with G-buffer. Septins were also polymerized in the absence of actin by overnight dialysis against a low-salt buffer (50 mM Tris-HCl pH 8, 50 mM KCl, 5 mM MgCl_2_, 1 mM DTT) at 4°C, then loaded into PLL-PEG-passivated flow channels in the presence of 1 mM Trolox, 2 mM PCA, 0.1 μM PCD and 0.1% w/v methylcellulose, and sealed as described above for observation. Septins were used at 20% or 100% GFP molar ratio, yielding similar results. Septin polymerization assays were repeated at least five times, using at least two different preps from each nonfluorescent and fluorescent hexamers, 8-mers-9_i1 and 8-mers-9_i3, and one prep from fluorescent 8-mers-9_i5, yielding similar results.

Actin-septin samples with mammalian septin hexamers (Fig. S3 C) were prepared as above with the difference that septins were nonfluorescent, and fluorescent actin was Alexa Fluor 488-conjugated G-actin (10% molar ratio) as described previously (Mavrakis et al., 2014; Mavrakis et al., 2016). G-actin polymerization in this case occurred in the presence of nonfluorescent phalloidin.

The sources and identifiers for proteins, materials and chemicals are as follows. Rabbit skeletal muscle G-actin from Cytoskeleton, Inc. (AKL99). Alexa Fluor 568-phalloidin from Thermo Scientific (A12380). Nonfluorescent phalloidin from Sigma (P2141). Methylcellulose from Sigma (M0512). Trolox from Sigma (238813). Protocatechuic acid from Sigma (03930590). Protocatechuate 3,4-dioxygenase from Sigma (P8279). 20K MWCO Slide-A-Lyzer MINI dialysis devices from Thermo Scientific (69590).

### Fluorescence microscope image acquisition and processing

Samples were imaged on an optical setup employing a confocal spinning disk unit (CSU-X1-M1 from Yokogawa) connected to the side-port of a Perfect Focus System-equipped inverted microscope (Eclipse Ti2-E from Nikon Instruments), using a Nikon Plan Apo ×100/1.45 NA oil immersion objective lens, 488- and 561-nm Sapphire laser lines (Coherent) and an iXon Ultra 888 EMCCD camera (1024×1024 pixels, 13×13 μm pixel size, Andor, Oxford Instruments), resulting in an image pixel size of 65 nm. Images were acquired with an exposure time of 0.1 s. Time-lapse sequences were acquired with a time interval of 0.5 s for a duration of 15 s. Actin filaments and actin-septin bundles were imaged close to the surface. Septin filament bundles were also found at the surface, but the extensive clusters of interconnected human septin filament bundles were observed floating in the bulk of the flow channels. To capture such clusters, z-stacks were acquired over 10-50 μm using a Δz interval of 0.5 μm. The images shown correspond to octamers-9_i1 polymerized with 20% GFP-septins (Fig. 3 A), octamers-9_i3 polymerized with 100% GFP-septins (Fig. 3 B), octamers-9_i5 polymerized with 100% GFP-septins (Fig. 3 C), hexamers polymerized with 100% GFP-septins (Fig. S 2 A), and *Drosophila* hexamers polymerized with 20% GFP-septins (Fig. S 2 B). All examples shown depict polymerization upon dilution into low salt apart from Fig. S 2 A (left panel) which shows polymerization upon dialysis into low salt.

Images were processed with the open-source image processing software ImageJ/Fiji. Images of actin filaments and actin-septin bundles are from single planes. Images of septin filament bundles are from maximum-intensity z projections except for *Drosophila* septins, for which single planes are shown given that their bundles were typically found primarily at the surface. The contrast of all images shown was adjusted post-acquisition so that both dim and bright structures are visible. To saturate the signal in the actin bundles and make the weaker-intensity signal of single/thinner actin filaments visible, the contrast was enhanced on purpose (images labeled “contrast enhancement” in Fig. 4 C-E and Fig. S3 A-B). All images shown use an inverted grayscale, with bright signals appearing black in a white background.

Actin-septin samples with mammalian septin hexamers (Fig. S3 C) were imaged with a Nikon Apo TIRF ×100/1.49 NA oil immersion objective lens mounted on an Eclipse Ti microscope (Nikon Instruments) using a 491 nm laser line and imaged with a QuantEM 512SC EMCCD camera (Photometrics). Images were acquired with an exposure time of 0.1 s.

### Transmission electron microscopy

#### Negative stain electron microscopy

4 μL of sample at final septin concentrations of 0.01-0.02 mg/mL (∼25-50 nM) for high salt conditions (50 mM Tris-HCl pH 8, 300 mM KCl, 2 mM MgCl_2_) or 0.05-0.1 mg/mL (∼125-250 nM) for low salt conditions (50 mM Tris-HCl pH 8, 50 mM KCl, 2 mM MgCl_2_) were adsorbed for 30 s (for high salt conditions) to 1 h in a humid chamber (for low salt conditions) on a glow-discharged carbon-coated grid (Electron Microscopy Sciences, CF300-CU). For low salt conditions, septins were polymerized by dilution into low-salt buffer and incubated for 1 h to overnight at room temperature before grid adsorption. In the case of GFP-labeled septins, septins were polymerized without mixing with nonfluorescent ones. The grids were rinsed and negatively stained for 1 min using 1% w/v uranyl formate. Images for the qualitative examination of the morphology of septin assemblies were collected using a Tecnai Spirit microscope (Thermo Scientific, FEI) operated at an acceleration voltage of 80kV and equipped with a Quemesa camera (Olympus). In addition to the EM experiments described above which were performed at I. Curie, EM was also performed at TU Delft following a similar protocol. Septins were polymerized by dilution into a low-salt buffer (25 mM Tris-HCl pH 7.4, 50 mM KCl, 2.5 mM MgCl_2_, 1 mM DTT) at a final septin concentration of 1 μM for 1 h. The solution was then adsorbed to a glow discharged grid for 1 min, rinsed, negatively stained with 2% w/v uranyl acetate for 30 s and air dried. Images were collected with a JEM-1400plus TEM microscope (JEOL) operated at 120kV equipped with 4k X 4k F416 CMOS camera (TVIPS). Septin filament bundle length and width measurements (Fig. S2 E) were made with the line tool in ImageJ/Fiji, and boxplots generated in Matlab. Electron microscopy was performed with at least two different preps from each nonfluorescent and fluorescent hexamers, 8-mers-9_i1 and 8-mers-9_i3. The images shown correspond to nonfluorescent octamers-9_i1 (Fig. 3 D, i-ii) and octamers-9_i3 (Fig. 3 E), SEPT2-msfGFP octamers-9_i1 (Fig. 3 D, iii-v), SEPT2-msfGFP hexamers (Fig. S 2 C), and nonfluorescent mammalian hexamers (Fig. S 2 D).

#### Two-dimensional image processing for single-particle EM images

Images for single-particle analysis were collected using a Lab6 G2 Tecnai microscope (Thermo Scientific) operated at an acceleration voltage of 200 kV. Images were acquired with a 4k X 4k F416 CMOS camera (TVIPS) in an automated manner using the EMTools software suite (TVIPS) with a pixel size of 2.13 Å and an electron dose of about 15 electrons/Å^2^. 2D processing was carried out on septin rods incubated in high salt conditions (50 mM Tris-HCl pH 8, 300 mM KCl, 2 mM MgCl_2_) to determine the integrity of the complexes and pinpoint the arrangement of septin subunits within the complex. About 100 images were collected for each septin complex for image processing. Individual particles (septin rods) were hand-picked from the images using the boxer tool from the EMAN software suite (Ludtke et al., 1999). About 20-30 particles (203×203 pixel boxes) were extracted per image. Subsequent processing was carried out using SPIDER (Frank et al., 1996). After normalization of the particles, a non-biased reference-free algorithm was used to generate 20 classes. Those classes were further used as references to pursue 2D multivariate statistical analysis. Multi-reference alignment followed by hierarchical classification involving principal component analysis was thereafter carried out to generate classes containing 50-100 particles. Each of the classes are representative of specific features within a given sample. This processing enabled us to quantify the distribution of particles in each dataset regarding the dimension of the rods as well as the presence of an additional electron density (GFP-tag or antibody). For mammalian octamers-9_i3 (Fig. 2 B), 4000 particles were selected with a distribution of 50% octamers, 23.7% heptamers, 23.5 % hexamers, 1.4% pentamers and 1.4% tetramers. For human SEPT2-msfGFP octamers-9_i1 (Fig. 2 C), 3266 particles were picked with a distribution of 57.9% octamers, 32.1% heptamers and 10.1% hexamers. An additional density towards the ends of the rods was pinpointed for 46.2% of the particles. For human SEPT2-msfGFP hexamers (Fig. 2 D), 2976 particles were selected with a distribution of 97.7% hexamers and 2.3% pentamers. An additional density towards the ends of the rods could be pinpointed for 53.6% of the particles. For human octamers-9_i1 incubated with SEPT9 antibodies (Fig. 2 C), 2352 particles were picked with a distribution of 13.7% octamers, 72.1% heptamers and 14.2% hexamers. 52.6% of the dataset exhibited an additional density towards the center of the rod that could be related to the presence of the antibody. Septin octamers were incubated with a rabbit antibody against the very C-terminus of SEPT9 (Pizarro-Cerda et al., 2002) for 30 min prior to grid adsorption and staining.

## Acknowledgements

We thank Josette Perrier and Cendrine Nicoletti (iSm2, Marseille, France) for generously hosting protein production and purification experiments; Christophe Romier (IGBMC, Strasbourg, France) and Jean-Denis Pedelacq (IPBS, Toulouse, France) for advise on protein purification; Jeffrey den Haan (TU Delft, The Netherlands) for help with protein purification; Tamara Advedissian and Arnaud Echard (I. Pasteur, Paris, France) for the gift of anti-SEPT9 antibody; Cristel Chandre (I2M, Marseille, France) for help with Matlab code; Caio Vaz Rimoli, Louwrens van Dellen and Sophie Brasselet (I. Fresnel, Marseille, France) for the development of the spinning disk optical setup and image acquisition software. This research received funding from the Agence Nationale pour la Recherche (ANR grant ANR-17-CE13-0014; SEPTIMORF), the Fondation ARC pour la recherche sur le cancer (grant PJA 20151203182), the Fondation pour la Recherche Medicale (FRM grant ING20150531962) and the Cancéropôle PACA and INCa. This project has received funding from the European Research Council (ERC) under the European Union’s Horizon 2020 research and innovation programme (grant agreement No 723241). This work was further financially supported by the Netherlands Organization for Scientific Research (NWO/OCW) through a VIDI grant (project number: 680-47-233) and the ‘BaSyC— Building a Synthetic Cell’ Gravitation grant (024.003.019), and from two PHC Van Gogh grants (no. 25005UA and no. 28879SJ, ministères des Affaires étrangères et de l’Enseignement supérieur et de la Recherche). Proteomic analyses were done using the mass spectrometry facility of Marseille Proteomics (marseille-proteomique.univ-amu.fr) supported by IBISA (Infrastructures Biologie Santé et Agronomie), Plateforme Technologique Aix-Marseille, the Cancéropôle PACA, the Provence-Alpes-Côte d’Azur Région, the Institut Paoli-Calmettes and the Centre de Recherche en Cancérologie de Marseille, the Fonds Européen de Developpement Régional and Plan Cancer. We further acknowledge the Cell and Tissue Imaging platform (PICT IBiSA, Institut Curie) supported by France-BioImaging (ANR10-INBS-04).

The authors declare no competing financial interests.

## Author contributions

F. Iv, A. Llewellyn, M. Belhabib, L. Ramond, A. Di Cicco, K. Nakazawa: investigation; C. Silva Martins, G. Castro-Linares, C. Taveneau, F-C Tsai, P. Barbier, P. Verdier-Pinard, L. Camoin, S. Audebert, investigation, writing – review & editing; R. Vincentelli, supervision, resources; J. Wenger, funding acquisition, supervision, writing – review & editing; S. Cabantous, supervision, writing – review & editing; G. H. Koenderink, conceptualization, funding acquisition, supervision, writing – review & editing; A. Bertin, conceptualization, supervision, investigation, writing – review & editing; M. Mavrakis, conceptualization, methodology, funding acquisition, project administration, supervision, investigation, writing – original draft, writing – review & editing

