## Supplemental material for "Insights into animal septins using recombinant human septin octamers with distinct SEPT9 isoforms"

### Online supplemental material

Fig. S1 contains details on the plasmids and septin sequences used for the isolation of recombinant human septin hexamers and octamers. Fig. S1 further contains additional data on the biochemical and biophysical characterization of septin complexes that relate to Fig. 1. Fig. S2 shows fluorescence and electron microscopy data on septin assembly from recombinant human, mammalian and *Drosophila* septin hexamers, and relates to Fig. 3. Fig. S3 shows fluorescence microscopy data on septin-actin reconstitution using recombinant human, mammalian and *Drosophila* septin hexamers, and relates to Fig. 4.

Figure S1. **Isolation and characterization of recombinant human septin octamers containing distinct SEPT9 isoforms. (A)** Schematic of the two sets of plasmids used for the co-expression of septins for isolating recombinant human SEPT2-, SEPT6-, SEPT7-containing hexamers (left) and recombinant human SEPT2-, SEPT6-, SEPT7-, SEPT9_i-containing octamers (right) from bacteria. SpeI and XbaI restriction sites used for subcloning are indicated (see Materials and methods for details). The end subunit (SEPT2) contains an N-terminal TEV-cleavable $His_6$-tag (depicted as H in the cartoons of the septin complexes), while the central subunit (SEPT7 in hexamers, or SEPT9_i in octamers) contains a C-terminal TEV-cleavable Strep-tag II. The plasmids for the production of fluorescent septin complexes used in this study differ only in that the gene coding for SEPT2 has been replaced by the one encoding SEPT2-msfGFP. **(B)** Primary sequences of the N- and C-termini of tagged septins used in the purification schemes in this study. $His_6$-tag, Strep-tag-II, and TEV cleavage site sequences are highlighted in light orange. Black arrowheads indicate the position of the TEV cleavage site. Asterisks point to the last amino acid of the respective septin sequence. The underlined amino acids in the C-terminus of SEPT2 are three out of the five residues that differ among the mouse and human homologs. The underlined stretch of N-terminal residues in SEPT7 was missing in previously reported plasmids (see Materials and methods for details). SEPT9 long isoform-specific sequences are highlighted in pink and cyan (see Fig. 1 B). **(C-E)** SDS-PAGE analysis of the purification of mammalian (C and D) and human (E) SEPT2-, SEPT6-, SEPT7-containing hexamers. Coomassie-stained gels show fractions from the total lysate (T), supernatant (S/N), flow-through (F/T), wash (W), eluate (E), and after concentration (C), using a two-tag purification scheme employing either a nickel affinity step followed by a Strep-Tactin affinity step (C), or a Strep-Tactin affinity step followed by a nickel affinity step (D and E). Molecular weight markers are shown on the left of each gel. The identification of bands is based on mass spectrometry analysis. The asterisk in (C) points to putative $His_6$-tagged SEPT2 homodimers that are removed in the Strep-tag affinity step. (**F**) Purified recombinant nonfluorescent and fluorescent (SEPT2-msfGFP) 6mer, 8mer-9_i1 and 8mer-9_i3 were analyzed by SDS-PAGE, followed by Western blot (WB) with antibodies against SEPT2, SEPT6, SEPT7, and SEPT9, as indicated at the bottom of each gel (see Materials and methods for details). Molecular weight markers are shown for the first gel; the same markers were used in all gels. All septins were intact, the long N-terminal extension of SEPT9 being most sensitive to proteolysis (the asterisk points to a degradation product for SEPT9). See Materials and methods for the theoretical and apparent molecular masses. **(G-H)** Examples of mass spectrometry analysis of recombinant 6mer, 8mer-9_i1 and 8mer-9_i3 preps. Calculations of the mol fractions of septins and contaminants in the respective protein preps are shown (G) using the Top3 quantitation approach (see Materials and methods for details). The obtained mol fractions of septins, compared with the theoretical ones in 6mer (33%) and 8mer (25%), point to the isolation of stoichiometric 6mers and 8mers. Examples of tryptic peptide coverage for individual septins in recombinant 6mer, 8mer-9_i1 and 8mer-9_i3 preps (H), supporting that the isolated septin complexes are intact. **(I)** Models for octamers without coiled-coils, or with coiled-coils at 90° with respect to the $\alpha 6$ helix, pointing to the same or to opposite directions as shown on the right, were used to calculate their theoretical sedimentation coefficients (see Materials and methods for details). The absence of coiled-coils altogether is predicted to make the complexes sediment faster by ~ 1.7 S units. Coiled-coils lying on the same side tend to make complexes more compact and thus slightly accelerate sedimentation by ~ 0.1-0.2 S units., whereas coiled-coils on opposite sides are predicted to slow down sedimentation by ~ 0.8 S units.

Figure S2. ***In vitro* reconstitution of septin polymerization in solution using recombinant animal septin hexamers. (A-B)** Representative spinning disk fluorescence images of higher-order filament assemblies upon polymerization of human SEPT2-, SEPT6-, SEPT7-containing hexamers (A) and *Drosophila* DSep1-, DSep2-, Peanut-containing hexamers (B) after dilution into low salt conditions (50 mM KCl) at the indicated final concentration. Two different *Drosophila* hexamers are shown: hexamers labeled with mEGFP-DSep2 (left panel in B) and hexamers labeled with DSep1-msfGFP (right panel in B). Two examples are shown for each type of hexamer. *Drosophila* hexamers organize in straight needle-like bundles, in line with previous reports (Mavrakis et al., 2014; Mavrakis et al., 2016). The freehand line preceding the G domain of Peanut in the hexamer cartoon above the images depicts its large N-terminal extension. Images in (A) are maximum-intensity projections. All images shown use an inverted grayscale. **(C)** Negative-stain EM images of higher-order filament assemblies upon polymerization of human 6mer at 0.2 $\mu$M and at low salt (50 mM KCl). The insets show magnifications of selected regions of interest (dashed rectangles in red), and highlight single septin filaments (blue arrowheads), paired septin filaments (orange arrowheads), and splayed filament bundles (i, iii). **(D)** Negative-stain EM of higher-order filament assemblies upon polymerization of mouse SEPT2-, human SEPT6-, human SEPT7$\Delta$N19-containing hexamers at low salt (50 mM KCl) and at 1 $\mu$M (i, ii) or 0.5 $\mu$M (iii). The insets show magnifications of selected regions of interest (dashed rectangles in red), and highlight single septin filaments (blue arrowheads), paired septin filaments (orange arrowheads), and splayed filament bundles (ii). **(E)** Box plots showing the distribution of septin filament bundle lengths (left), septin filament bundle widths (middle) and septin filament widths within bundles (right), measured from electron micrographs, and comparing human 6mer- (red-filled circles) and 8mer-9_i1 (blue-filled circles) filament assemblies. The data points are plotted on top of the respective box plots. On each box, the central mark indicates the median, and the bottom and top edges of the box indicate the 25th and 75th percentiles, respectively. The whiskers extend to the most extreme data points not considered outliers, and the outliers are plotted individually using the `'x'` symbol. The number of measurements in each box plot, ordered from left to right, is n = 58, 83, 229, 69, 30, 28. The respective median values are 1.9 μm, 1.3 μm, 78 nm, 67 nm, 4.6 nm, and 3.9 nm.

Figure S3. ***In vitro* reconstitution of actin filament cross-linking by recombinant animal septin hexamers. (A-C)** Representative spinning disk fluorescence images of reconstituted actin filaments, polymerizing in the presence of human SEPT2-, SEPT6-, SEPT7-containing hexamers (A), *Drosophila* DSep1-, DSep2-, Peanut-containing hexamers (B), and mouse SEPT2-, human SEPT6-, human SEPT7ΔN19-containing hexamers (C), prepared as in Fig.4 C-D. (A-B) Actin filaments are visualized with AlexaFluor568-conjugated phalloidin, and septins with SEPT2-msfGFP (human) or DSep1-msfGFP (*Drosophila*). Two examples of large fields of view are shown for each, depicting the similar cross-linking of actin filaments into actin filament bundles in the presence of both types of hexamers; only actin labeling is shown. Insets on the right side of each panel show higher magnifications of selected regions of interest on the left (dashed squares in red). Two regions of interest (i, ii) are shown in each case, depicting both the actin (top row) and septin (bottom row) signals. For each inset, actin and septin signals are shown in duplicates: the first set shows the raw signals without any saturation, whereas the second set, adjacent to the first one, shows both actin and septin signals after deliberate contrast enhancement. The contrast-enhanced images in the actin channel saturate the actin bundles, while bringing out weaker-intensity single actin filaments (black arrowheads). The respective contrast-enhanced images in the septin channel show the presence of septins in actin bundles, but their absence from single actin filaments. Scale bars in all large fields of views, 10 μm. Scale bars in all insets, 5 μm. (C) Actin filaments are visualized with Alexa Fluor 488-G-actin and septins are nonfluorescent. Three examples of large fields of view are shown, depicting the similar cross-linking of actin filaments into actin filament bundles. Scale bars in all large fields of views, 10 μm. All images shown use an inverted grayscale.

Video 1. **Polymerization of recombinant human septin octamers-9i3.** Optical sectioning (z-stack with a Δz interval of 0.5 μm) in the bulk of a flow channel depicting SEPT2-msfGFP human septin octamer-9_i3 polymerized at 0.3 μM (100% GFP-septins) by dilution into low-salt (50 mM KCl) buffer. Spinning disk fluorescence images displayed at 5 frames per second. Related to Fig. 3 B.

Video 2. **Reconstitution of single actin filaments.** Time-lapse sequence ($\Delta$t interval of 0.5 s) at the surface of a PLL-PEG passivated glass coverslip showing single fluctuating actin filaments at 1 $\mu$M. G-actin was polymerized in the presence of Alexa Fluor 568-phalloidin. Spinning disk fluorescence images displayed at 5 frames per second. A still image from this time lapse sequence is shown in Fig. 4A.

Video 3. **Actin filament cross-linking by recombinant human septin octamers-9i1.** Time-lapse sequence ($\Delta$t interval of 0.5 s) at the surface of a PLL-PEG passivated glass coverslip showing cross-linked actin filaments (at 1 $\mu$M) in the presence of SEPT2-msfGFP human septin octamer-9_i1 at 0.3 $\mu$M (20% GFP-septins). G-actin was polymerized in the presence of Alexa Fluor 568-phalloidin. The actin channel is shown. Spinning disk fluorescence images, using an inverted grayscale, are displayed at 5 frames per second. A still image from this time lapse sequence is shown in Fig. 4C.

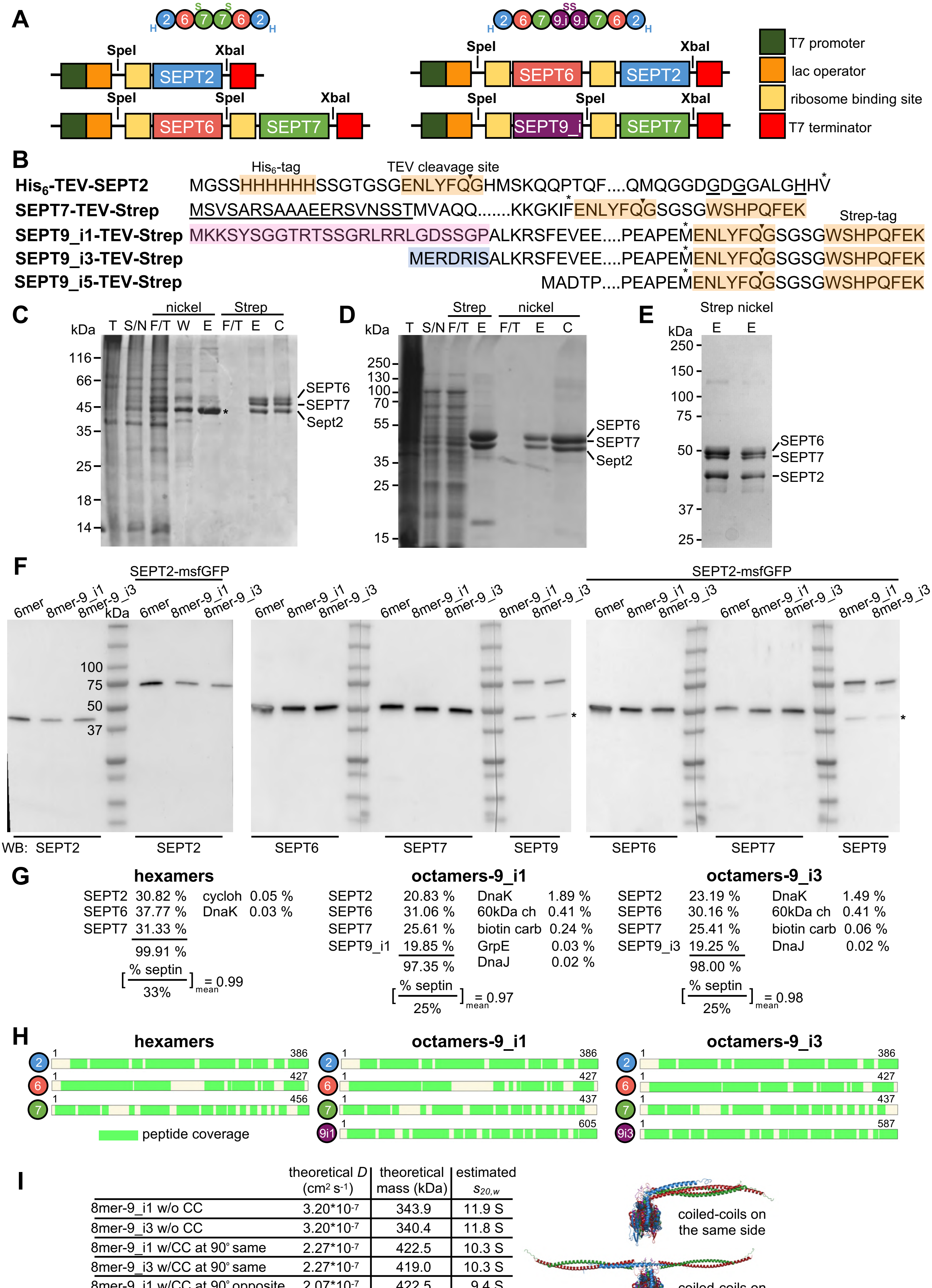


**Figure S1**
**Iv et al 2020**

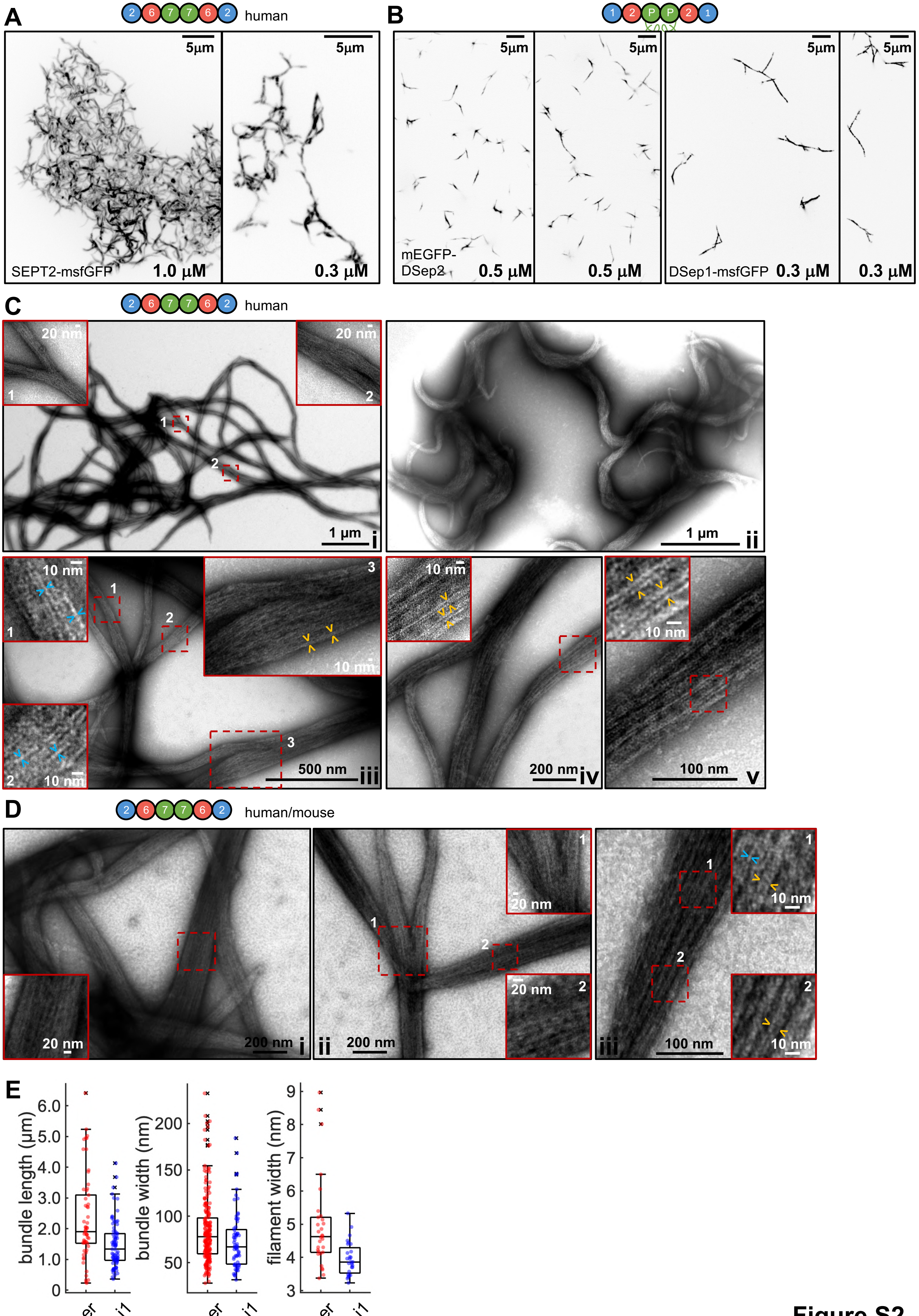


Figure S2
Iv et al 2020

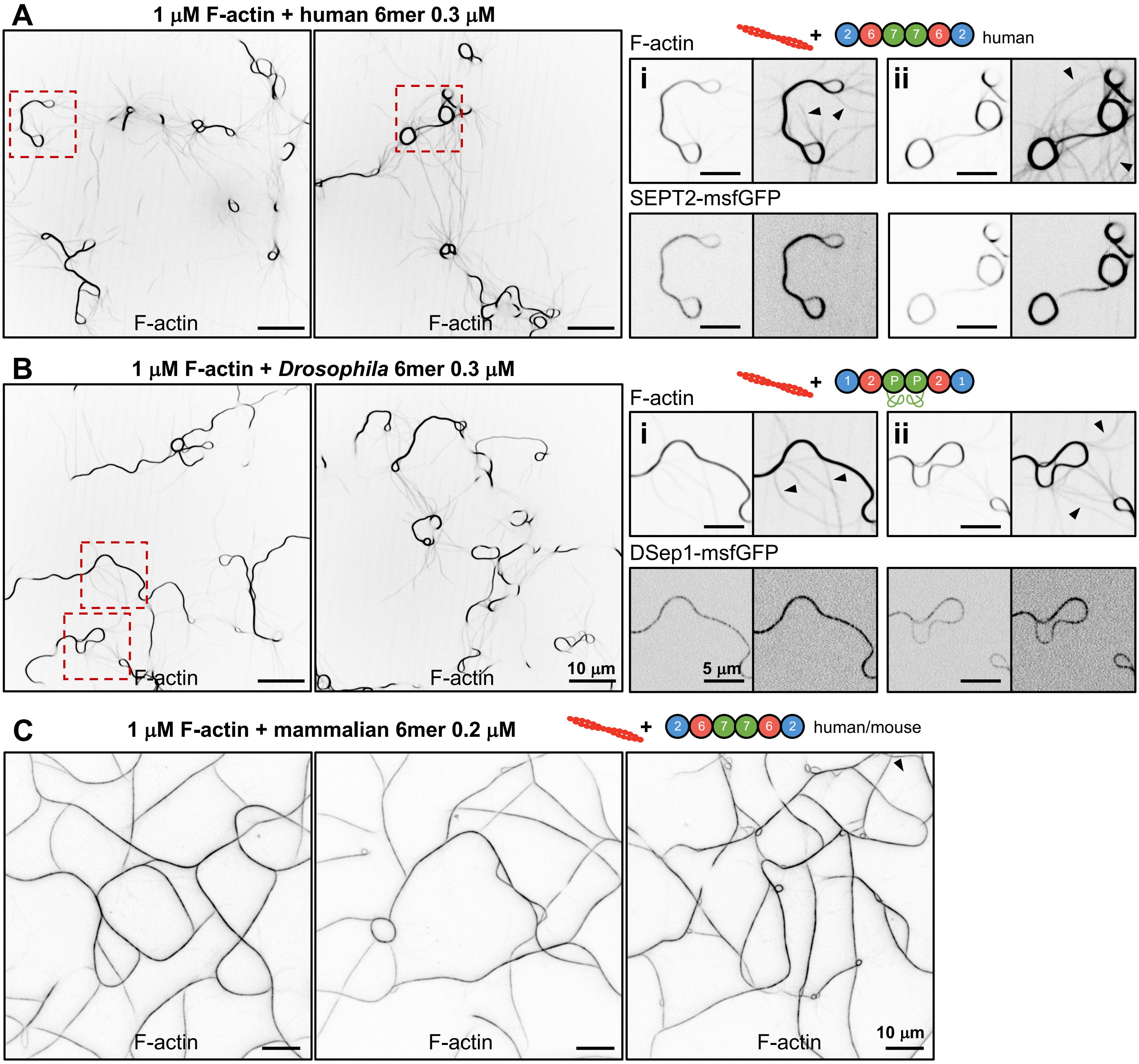


**Figure S3**
**lv et al 2020**
